# Thermoregulation network governing virulence of a critical human fungal pathogen

**DOI:** 10.1101/2025.08.18.670910

**Authors:** Vikas Yadav, Joseph Heitman

## Abstract

*Cryptococcus neoformans* is an environmental fungal pathogen that causes meningoencephalitis in humans, requiring thermal adaptation to human body temperature. We employed several orthogonal complementary approaches to elucidate molecular mechanisms of calcineurin signaling, which is essential for thermotolerance of many fungal species, and in so doing, delineated a thermoregulatory network. First, genetic suppressors for loss of calcineurin activity identified mutations in a kinase, Yak1, as the primary suppression mechanisms. Second, the development and utilization of the proximity labeling tool TurboID identified novel subcellular interactions of calcineurin during thermal stress. Third, investigations employing phosphoproteome, RNA-sequencing, and Ribo-sequencing revealed a major role for calcineurin in controlling translation initiation machinery during thermal stress adaptation. Fourth, truncation alleles revealed domain-specific roles of calcineurin catalytic A subunit in thermotolerance, meiosis, and virulence. Combined, this study presents a comprehensive analysis of thermotolerance-governing networks and mechanisms in a fungal pathogen of global impact.

## Introduction

Fungi pose a growing threat in the increasingly warming world, endangering food security, biodiversity, and human health. The risk to human health is increasing due to the growing population undergoing immunosuppressive medical treatments. In the absence of a robust immune system, human body temperature (37°C) is one of the major defense systems because most fungi are unable to grow at 37°C. The majority of fungi thrive in cooler environments, but increasing global temperatures are proposed to drive changes that will prime more fungal species to grow at higher temperatures and threaten human health (Casadevall *et al*, 2019; Cordero *et al*, 2023; Ramos Irizarry *et al*, 2025). Thus, it is critical to understand the mechanisms by which fungi adapt to higher temperatures, specifically 37°C, and even more so for fungal species that already cause infections.

*Cryptococcus neoformans* is an environmental fungus that primarily infects both immunocompromised and immunocompetent hosts and is responsible for >147,000 deaths globally with a 75% mortality rate (Denning, 2024). Due to its significance, it is listed as the top critical priority fungal pathogen by the World Health Organization (W.H.O., 2022). Thermotolerance is known to be a crucial virulence factor for *C. neoformans,* and mutants unable to grow at 37°C fail to cause infections (Kwon-Chung *et al*, 2014; Odom *et al*, 1997; Perfect, 2006). Calcium-calcineurin signaling is a highly conserved pathway across eukaryotes and is required for growth at 37°C and pathogenesis in *C. neoformans* (Odom *et al*., 1997). The central component in this signaling cascade is calcineurin, a phosphatase consisting of the catalytic subunit, Cna1, and the regulatory subunit, Cnb1. In humans, calcineurin plays crucial roles in T-cell activation during immune function, and also in neurological and cardiovascular disease (Creamer, 2020; Klee *et al*, 1979; Park *et al*, 2019; Roy & Cyert, 2020; Ulengin-Talkish & Cyert, 2023). Due to its role in T-cell activation, the FDA-approved calcineurin inhibitors FK506 and cyclosporin A (CsA) are gold standard immunosuppressive therapies widely applied to prevent graft rejection in organ and tissue transplant recipients (Hemenway & Heitman, 1999; Ho *et al*, 1996).

In fungi, calcineurin signaling is essential for virulence and regulates morphological transitions and stress responses (Park *et al*., 2019; Yadav & Heitman, 2023). In *C. neoformans*, in addition to its essential role in thermotolerance, calcineurin is also required for sexual reproduction by promoting the yeast-to-hyphal transition after mating (Cruz *et al*, 2001). Previous studies identified the conserved calcineurin-dependent downstream signaling transcription factor Crz1 and showed that calcineurin localizes to P-bodies/stress granules, where it plays a role in small-interfering RNA production mediating RNAi (Chow *et al*, 2017; Kojima *et al*, 2006; Kozubowski *et al*, 2011a; Park *et al*, 2016; Yadav *et al*, 2024). However, the regulatory roles of calcineurin at a sub-cellular level as well as its interactions with other cellular networks remained uncharacterized.

Here, we employed several parallel approaches to define the calcineurin thermoregulatory network in *C. neoformans*. Through two genetic screens, one based on spontaneous mutations and another with a targeted protein kinase deletion library, we identified Yak1 as the primary kinase that acts antagonistically to calcineurin at 37°C. We developed and optimized a proximity ligation-based approach, TurboID, to identify novel proteins interacting with calcineurin, implicating roles for calcineurin in regulating microtubule dynamics, mRNA splicing, and the translation machinery, thereby impacting mitochondrial function. With a series of calcineurin A catalytic subunit truncation alleles, we delineated the roles of various domains present in Cna1 and identified domain-specific roles in meiosis and virulence. Overall, this study delineates a cellular map of the calcineurin signaling cascade and illustrates how these interactions form a functional network that governs thermal stress responses in a human fungal pathogen of critical importance.

## Results

### Genetic suppression analysis identified Yak1 as a major suppressor of calcineurin function

Calcineurin signaling is a major hub for stress responses in fungi and several subcellular functions such as cell wall organization, antifungal drug resistance, and morphological transitions (Juvvadi *et al*, 2017; Yadav & Heitman, 2023). To understand these functional interactions in *C. neoformans*, we conducted two parallel genetic screens. First, we isolated spontaneous genetic suppressor mutants capable of growing at 37°C in the absence of functional calcineurin (Figure S1A). We isolated 16 suppressor colonies from either 1) wild-type cells that grew in the presence of the calcineurin inhibitors FK506 and CsA at 37°C or 2) *cnb1*Δ mutant cells lacking the calcineurin B regulatory subunit that grew at 37°C. Suppressor mutants from both conditions partially suppressed growth under the restrictive conditions (Figure 1A-B and S1B).

**Figure 1.**
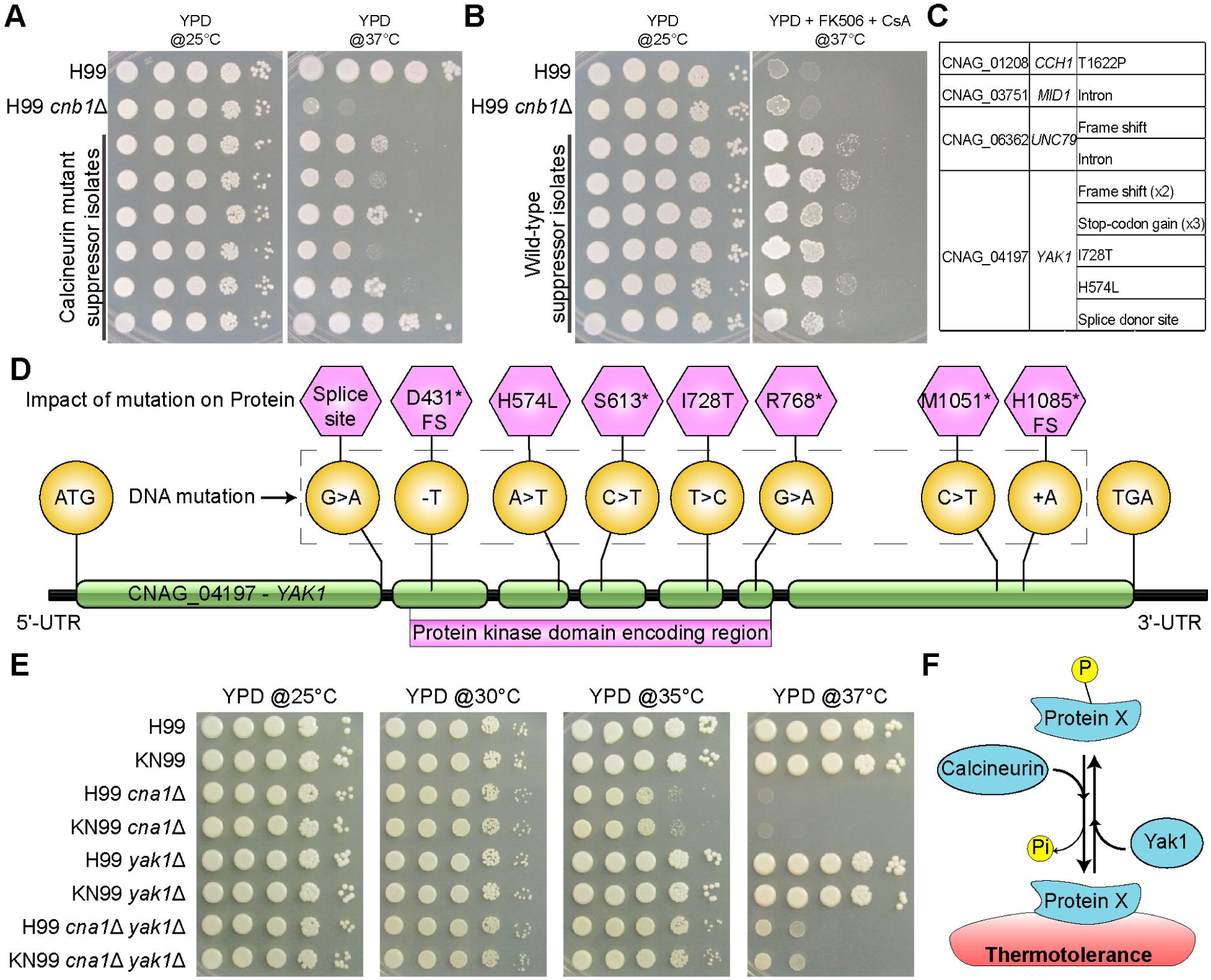
Genetic suppression analysis identified Yak1 as a major suppressor of calcineurin function. (A-B) Plate images showing the growth of isolated genetic suppressors as compared to the parent wild-type or *cnb1*Δ mutant strains in respective restrictive conditions. **(C)** A summary table with the list of identified mutations in each of the genes. **(D)** Structure of the gene (ORF number – *CNAG_04197*) encoding the protein kinase Yak1, which was identified to be mutated in 8 independent isolates. The nature of the DNA mutations (golden circles) and their impact on the protein (pink hexagons), as well as the start and stop codons, are depicted. * in the pink hexagons indicates gain of a stop codon, and FS refers to the frameshift nature of the mutation. Green color bars represent exons of the *YAK1* gene, and black lines mark untranslated regions (5’-UTR and 3’-UTR) as well as introns in between exons. The region of the gene that encodes the kinase domain is marked. **(E)** Plate images showing growth of wild-type strains (H99α and KN99**a**), respective calcineurin mutants (*cna1*Δ), *yak1* mutants (*yak1*Δ), as well as the double mutants (*cna1*Δ *yak1*Δ) at various temperatures. **(F)** A scheme depicting the relationship between calcineurin and Yak1 and the mechanism through which loss of Yak1 could potentially suppress the requirement for calcineurin function during high-temperature growth.

Whole genome sequence analysis of these mutants identified mutations in several genes as well as examples of segmental aneuploidy that could mediate suppression (Figure 1C and S1C). The aneuploid segment in chromosome 4 (isolates F2, F4, and H4), as well as two separate aneuploid regions in isolate H3, do not harbor any known genes implicated in suppressing calcineurin activity (Figure S1C). Interestingly, four of the isolates harbored a mutation in three components (Cch1, Mid1, and Unc79) of the NALCN calcium channel that was recently reported (Boucher *et al*, 2025). That study identified four subunits of the NALCN channel as a cluster of genes that exhibited higher growth in the presence of FK506 or CsA at 30°C from a genome-wide deletion library of *C. neoformans* (Figure S1D). Among our suppressor isolates, the highest number of mutations was identified in the gene encoding a kinase, Yak1 (8/16). Several of these mutations were frame shift or stop-codon gain mutations, suggesting that these are loss-of-function mutations (Figure 1C and 1D).

For the second genetic screen, we focused on a protein kinase deletion library consisting of 129 mutants (Lee *et al*, 2016). We hypothesized that a kinase might function antagonistically with calcineurin, regulating shared proteins to drive thermotolerance. This screen resulted in the identification of three kinase mutants that showed improved growth at 35°C and 37°C in the presence of the calcineurin inhibitors FK506 and CsA as compared to the wild type (Figure S1E). Among these, the *yak1*Δ mutant exhibited the most robust suppression of loss of calcineurin function, supporting the findings from the spontaneous mutant genetic screen. In addition to Yak1, this analysis resulted in the identification of two additional kinase mutants (*pan3*Δ and *pka1*Δ) that showed improved growth in the presence of FK506 and CsA as compared to the wild-type (Figure S1E). One of these, a protein kinase A catalytic subunit mutant, *pka1*Δ, exhibited growth restoration at 35°C but not at 37°C. The third kinase identified was Pan3, which harbors a pseudokinase domain and is involved in mRNA deadenylation, thus regulating RNA degradation (Christie *et al*, 2013; Schafer *et al*, 2014).

Both screens identified loss of Yak1 as a major suppressor for loss of calcineurin function during 37°C growth. To assess this finding further, we generated independent *cna1*Δ *yak1*Δ double deletion mutants, which showed growth restoration at 37°C as compared to the *cna1*Δ mutant alone (Figure 1E). Based on these findings, we hypothesize that calcineurin and Yak1 share certain substrates within the cell and control their function through dephosphorylation and phosphorylation, respectively (Figure 1F). To identify shared substrates between calcineurin and Yak1, we conducted phosphoproteome analysis of the *cna1*Δ and *yak1*Δ mutant strains at 25°C and 37°C (Figure 2A and 2B). Comparison of peptides that were hypophosphorylated (down) in the *yak1*Δ mutant and hyperphosphorylated (up) in the *cna1*Δ mutant identified 409 shared peptides, corresponding to 275 proteins (Figure 2C and Table S1). Gene ontology-based analysis of these genes revealed enrichment of microtubule-associated proteins, RNA-binding proteins, proteins involved in cellular trafficking, and other proteins involved in biological regulation (Figure 2D). We also detected Crz1, Vts1, and Lhp1 among the shared proteins, all of which have been previously shown to play an important role in calcineurin-mediated thermotolerance (Figure 2E) (Chow *et al*., 2017; Park *et al*., 2016).

**Figure 2.**
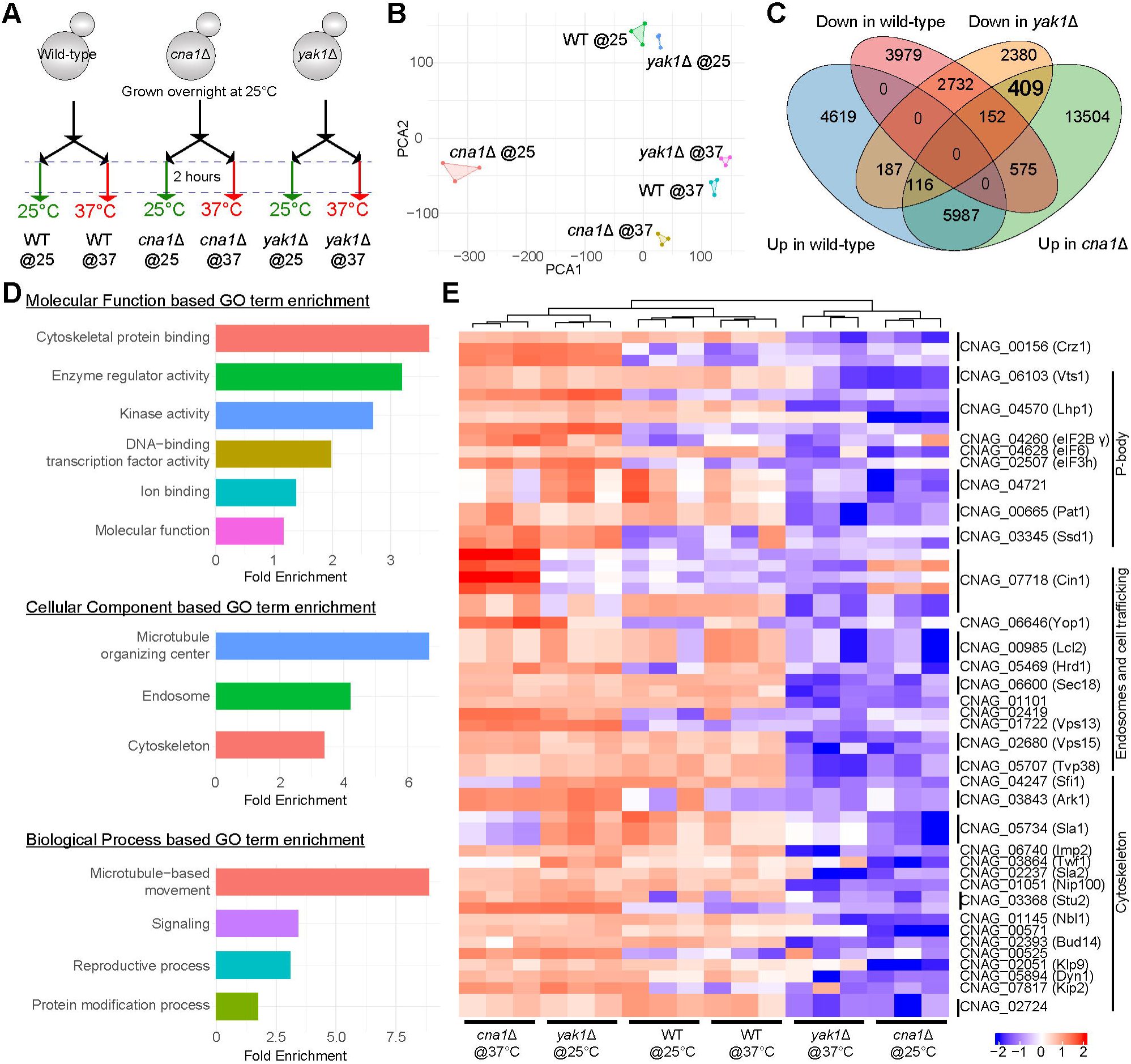
Phosphoproteomics analysis identified substrates shared between calcineurin and Yak1. **(A)** A cartoon scheme showing the phosphoproteome approach employed to identify phosphopeptides shared between calcineurin and Yak1. **(B)** A PCA plot showing the replicability among phosphoproteome samples. **(C)** A Venn diagram showing the overlap of peptide populations differentially regulated at 37°C for each strain as compared to 25°C. **(D)** Gene Ontology enrichment plots for the shared proteins corresponding to 490 peptides. **(E)** Heatmap showing the differential phosphorylation levels for peptides for selected proteins, including Crz1 and P-body proteins such as Vts1 and Lhp1.

Given that the *yak1*Δ mutant does not have a thermosensitive or thermoresistant phenotype, we think that it is responsible for phosphorylation of these proteins only at 37°C, and some of these phosphotargets may be redundantly regulated by other kinases. This was also supported by our analysis of phosphorylation states for two shared proteins, one of which is involved in actin depolymerization (CNAG_03864) and one encoding a dynein light chain subunit (CNAG_00525). Both of these proteins exhibited a lower overall population of phosphorylated protein upon calcineurin inhibition by FK506 (Figure S2). Both of these proteins harbor LxVP or PxIxIT calcineurin docking motifs, and it is possible that calcineurin directly dephosphorylates one or more sites that were identified by phosphoproteome analysis. However, calcineurin regulates several other kinases and phosphatases through dephosphorylation (described below), which in turn might be acting on these two proteins, resulting in less overall phosphorylation. Importantly, the observed decrease in phosphorylation of these proteins upon calcineurin inhibition suggests that their function may be impacted by calcineurin inhibition, albeit indirectly. Overall, these results reveal that Yak1’s antagonism to calcineurin function in governing thermotolerance occurs through modulation of multiple parallel cellular functions.

### TurboID proximity ligation revealed novel calcineurin interactions

To complement the genetic interaction analysis, we next characterized physical interactions of calcineurin in *C. neoformans*. Previous studies have identified calcineurin effectors via immunoprecipitation assays and phosphoproteome analysis and demonstrated roles for calcineurin in the secretory pathway as well as in P-bodies (Kozubowski *et al*, 2011b; Park *et al*., 2016). However, immunoprecipitation assays may not identify transient interactions, which are more likely to occur in signaling cascades, including enzyme-substrate interactions. To determine these interactions, we developed and optimized a proximity ligation assay, TurboID (Branon *et al*, 2018; Kalem *et al*, 2022), and applied it to Cna1 (Figure 3A and S3). A strain constitutively expressing TurboID (TID) served as a control to account for non-specific interactions that might arise from TurboID itself. Importantly, expression of TurboID alone for the control strain required optimization through the insertion of two introns within the gene, which stabilized the protein (Figure S3A). These strains exhibited robust growth at 37°C and underwent sexual reproduction when crossed with either the wild type or the *cna1*Δ mutant (Figure S3B-C). These results demonstrate that the Cna1-TID fusion protein is fully functional, and expression of TID alone does not impact growth at 37°C, providing functional validation. Notably, *C. neoformans* exhibits a high level of inherent biotinylation even in the wild-type strain (Figure S3D), which is essential for its viability as cells cease to grow after prolonged biotin starvation. Additionally, fungal cells can store biotin in their cells (Hasim *et al*, 2013; Naumova & Vorob’eva, 1983; Rogers & Lichstein, 1969), which can bind to streptavidin when released after cell lysis, and requires an additional step of purification to separate biotinylated proteins from free competing biotin.

**Figure 3.**
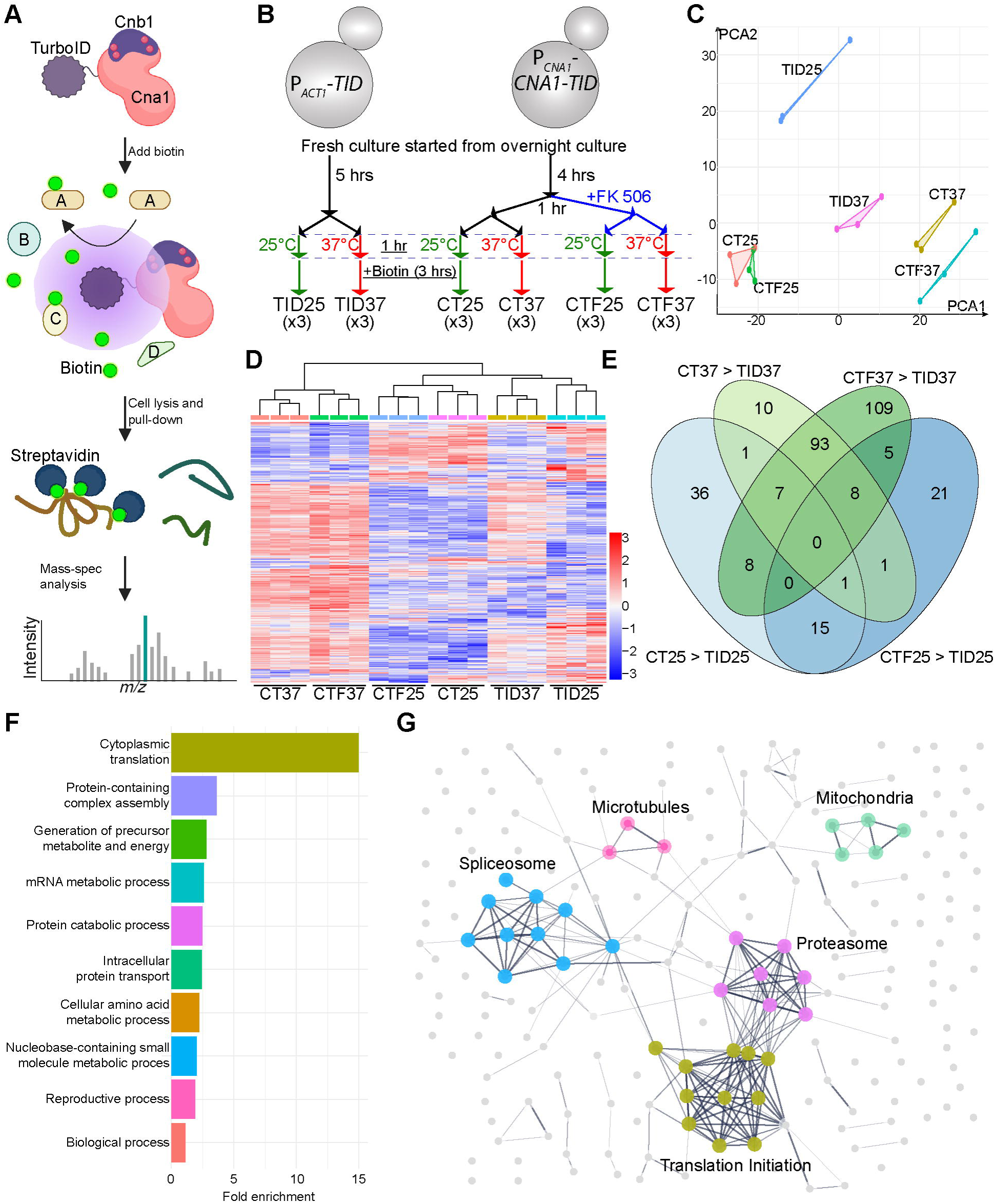
TurboID proximity ligation revealed novel calcineurin protein interactions. **(A)** A schematic depicting the principles of the TurboID assay to detect proteins interacting with calcineurin. **(B)** A cartoon showing the outline of the Cna1-TurboID (Cna1-TID) assay conducted with *C. neoformans* at two different temperatures, 25°C and 37°C. TID25 and TID37, TurboID control assays; CT25 and CT37, Cna1-TID assays in the absence of FK506; CTF25 and CTF37, Cna1-TID assays in the presence of FK506. **(C)** PCA analysis revealed a strong correlation between the three replicates for each condition. **(D)** Heatmap analysis after ANOVA multi-sample comparison revealed clustering of samples based on temperatures and provided additional support for replicate analysis. **(E)** Cna1-TID specific interactions were determined for each condition by pair-wise comparison of CT25 and CTF25 with TID25, and CT37 and CTF37 with TID37. The resulting sets of interacting proteins were analyzed and are presented as a Venn diagram. **(F)** Biological process-based gene ontology (GO) enrichment analysis of calcineurin interactions. **(G)** STRING network analysis showing the 5 highest interacting sets of proteins among the calcineurin interactome. The marked complexes are highlighted in colors, whereas the remaining proteins are depicted with grey circles.

Once optimized, we conducted biotinylation assays at both 25°C and 37°C, both in the presence and absence of the calcineurin inhibitor FK506 (Figure 3B, and S3D-E). PCA and heatmap analysis revealed a strong correlation between replicates (Figure 3C-D). Next, we defined the proteins interacting with Cna1 at both 25°C and 37°C after removing the proteins that were also present in the TurboID control. A comparison of these Cna1-specific proteins revealed several 37°C-specific interactions with a significant overlap between the group treated with FK506 and without treatment (Figure 3E). Furthermore, we identified a higher number of interacting proteins when FK506 was present, similar to our previous observation utilizing immunoprecipitation assays in the related species *C. deneoformans* (Yadav *et al*, 2025). For further analysis, we combined these two pools, resulting in a total of 212 37°C Cna1-interacting proteins (Table S2).

Gene Ontology (GO) analysis of these calcineurin interacting proteins revealed significant enrichment of proteins involved in several biological processes (Figure 3F). These encompassed proteins involved in intracellular transport as well as those localized to P-bodies, in accord with previous findings (Kozubowski *et al*., 2011b; Park *et al*., 2016). However, four of the most enriched biological processes included proteins from the translation initiation complex (GO: cytoplasmic translation), the spliceosome (GO: mRNA metabolic process), the proteasome machinery (GO: protein catabolic process), and mitochondria (GO: Generation of precursor metabolite and energy) (Figure 3F). STRING networking analysis further showed strong interactions of proteins involved in each of these categories (Figure 3G). While some of the translation initiation complex proteins have been reported to interact with calcineurin (Park *et al*., 2016), calcineurin’s interactions with the spliceosome and mitochondria were not previously characterized. Among 212 proteins, 61 have at least one peptide that is hyperphosphorylated in the calcineurin mutant at 37°C, with representation from each of these cellular processes (Figure 4A). Additionally, 103 of the 212 proteins harbor either or both of the calcineurin docking motifs, PxIxIT or LxVP (Figure S4A), suggesting that these could be direct substrates of calcineurin.

**Figure 4.**
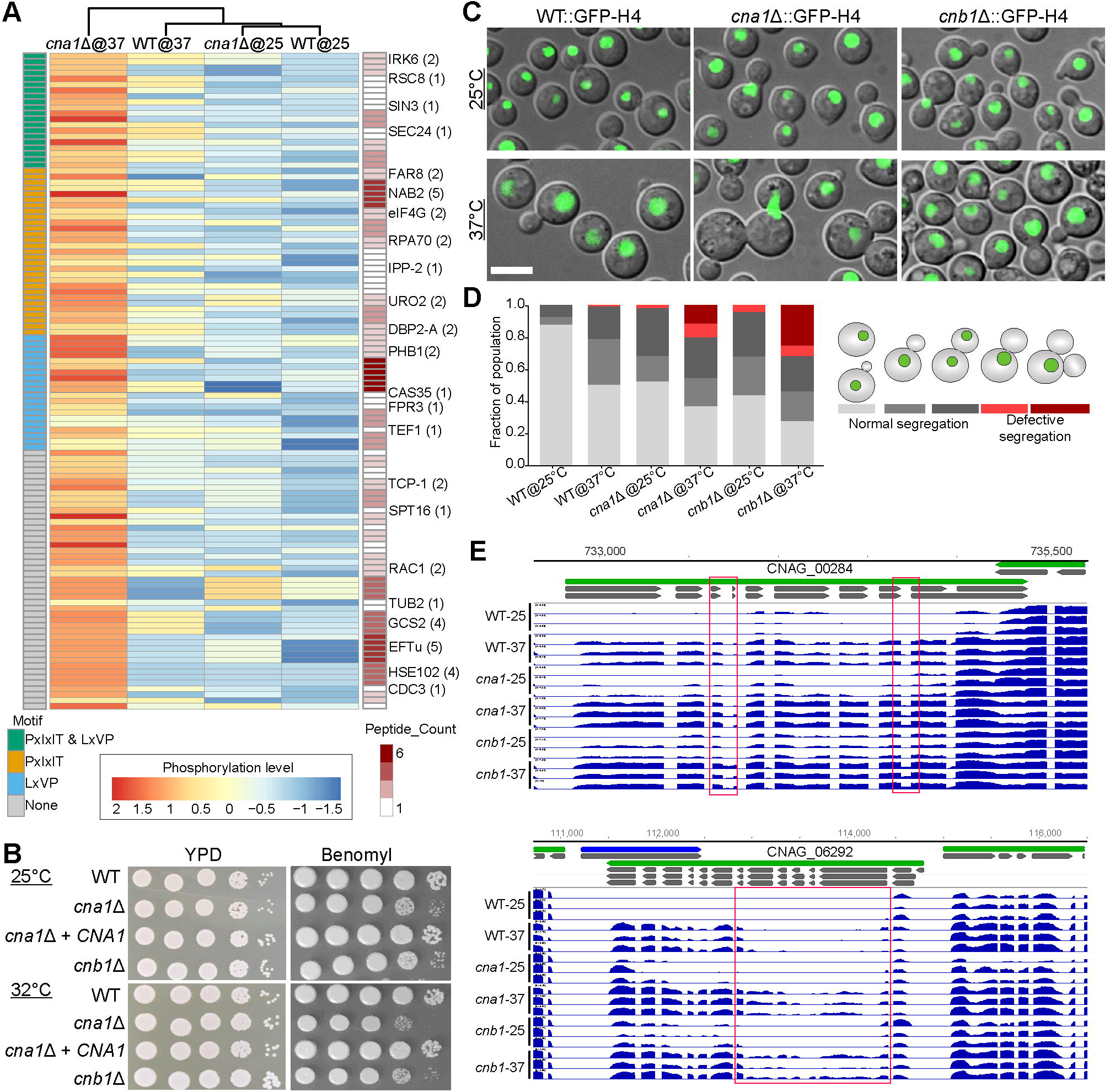
Calcineurin orchestrates several cellular processes. **(A)** A heatmap showing the Cna1-interacting proteins with at least one hyperphosphorylated peptide in the *cna1*Δ mutant at 37°C. The presence of PxIXIT and LxVP motifs, as well as the peptide count for each protein, are also presented. Only the proteins that are annotated are listed with their peptide count in parentheses. **(B)** Plate images showing sensitivity of calcineurin mutants to the microtubule-depolymerizing drug benomyl at 25°C or 32°C. **(C)** Fluorescence images showing nuclear segregation defects in calcineurin mutants at 37°C as compared to wild-type cells. Scale bar, 5 µm. **(D)** A graph presenting the cell population with defective nuclear segregation in the calcineurin mutants at 25°C and 37°C. **(E)** IGV plots showing examples of the altered splicing events in the calcineurin mutants at 37°C.

### Calcineurin regulates microtubule function

Several proteins involved in microtubule dynamics were identified by phosphoproteome analysis as substrates shared between calcineurin and Yak1, including the beta-tubulin subunit. The TurboID network analysis also identified interactions with all three proteins responsible for microtubule formation (one alpha-tubulin subunit and two beta-tubulin subunits). This prompted us to determine whether calcineurin plays a direct role in microtubule dynamics. We subjected calcineurin mutants to the microtubule-depolymerizing drug, benomyl, and found that calcineurin mutants are benomyl hypersensitive (Figure 4B). This suggests that calcineurin might regulate microtubule function in *C. neoformans*.

To directly study microtubule dynamics, we employed a strain expressing GFP-tubulin (Kozubowski *et al*, 2013) and found that microtubules formed longer and more stable strands in the absence of calcineurin at 25°C (Figure S4B). We were unable to conduct these assays at higher temperatures as the GFP-tubulin signal was unstable in calcineurin mutants at higher temperatures, further suggesting a functional interaction between calcineurin and microtubules. Microtubules have been shown to be involved in nuclear migration from mother to daughter cells during mitosis in this species (Kozubowski *et al*., 2013; Sutradhar *et al*, 2015). Thus, we determined whether calcineurin mutants exhibit nuclear migration defects at higher temperatures. Fluorescence imaging of GFP-tagged histone H4 localization in the wild-type and calcineurin mutants revealed a significant defect in the calcineurin mutants, where 10 to 20% of the cell population was multi-budded with only one nuclear signal at 37°C (Figure 4C-D and S4C). Furthermore, approximately 20% of calcineurin mutant cells with a daughter-to-mother ratio of 0.7 harbored a nucleus in the mother cell, whereas the nucleus was found to be divided in this cell population in wild-type cells. These results show that calcineurin plays an important role in regulating microtubule dynamics, resulting in delayed nuclear dynamics during mitosis.

### Calcineurin controls spliceosome function

One of the strongest Cna1-TurboID interactions observed was with components of the spliceosome, suggesting a potential role for calcineurin in splicing. Among the identified 10 interacting proteins, seven (CNAG_00111, CNAG_00147, CNAG_01091, CNAG_01733, CNAG_02901, CNAG_04069, CNAG_06585) were recently shown to be part of the splicing machinery in *C. neoformans* (Sales-Lee *et al*, 2021). This study, along with others, has shown that the splicing machinery involves several accessory factors that enable splicing of a diverse set of introns (Krishnan *et al*, 2024; Negi *et al*, 2025; Sales-Lee *et al*., 2021).

To directly determine the impact of calcineurin on RNA processing, we performed RNA-sequencing after 2 hours of heat stress at 37°C of the wild-type, *cna1*Δ, and *cnb1*Δ mutants (Figure S4D). The RNA sequencing data were then analyzed with the rMATS pipeline (Shen *et al*, 2014) to identify splicing variants with respect to wild-type cells grown at 25°C. Interestingly, the major impact on RNA splicing was found to be shared between wild-type and calcineurin mutants grown at 37°C, revealing that high temperature growth results in differential splicing that impacts a wide range of genes (Figure S4E). The majority of these genes encode for hypothetical proteins, and gene ontology enrichment analysis identified enrichment for proteins involved in the NAD biosynthetic process and mRNA metabolism pathway. These findings corroborate a previous study that identified a large impact of temperature on alternative transcription start sites resulting in gene expression changes and differential protein targeting (Dang *et al*, 2024). The deletion of calcineurin had a modest impact on splicing for a smaller subset of genes. Specifically, several intron retention as well as alternative splicing events were observed that were shared between both the *cna1*Δ and the *cnb1*Δ mutants (Figure 4E and Table S3). This analysis revealed that calcineurin regulates the function of the splicing machinery to a modest degree. Our results are also supported by a recent study showing a synergistic interaction between FK506 and the spliceosome inhibitor pladienolide B in *C. neoformans* (Love *et al*, 2025). It is possible that the impact on splicing was less pronounced with only 2 hours of heat stress in our assays. Alternatively, mRNAs with intron retention events may be detected by cellular machinery such as RNAi and degraded, resulting in lower levels of detection for such events.

### Calcineurin plays a major role in maintaining protein homeostasis

Among the Cna1-TurboID data, the most highly implicated cellular process was protein translation initiation. Additionally, several components of the proteasome machinery were detected, suggesting a global impact on protein homeostasis. Consistent with this interaction, deletion, or inhibition of calcineurin rendered cells hypersensitive to a proteasome inhibitor, bortezomib, at 32°C (Figure 5A). Among the translation initiation interactions, components of eukaryotic initiation factors (eIF) such as eIF2, eIF3, eIF4, and eIF5 were found to be associated with calcineurin. We performed parallel RNA-sequencing and Ribo-sequencing to identify the impact on protein translation in the *cna1*Δ mutant as compared to the wild-type at both 25°C and 37°C . Quality assessment for the Ribo-seq data revealed lower sequencing coverage and slightly larger ribosome-protected fragments (30-34 nt) as compared to a previous study in *C. neoformans* (Wallace *et al*, 2020), but is still within the expected range (Figure S5A). We think this could be due to differences in the sample preparation pipeline that involved treatment with harringtonine and cycloheximide along with slow cell lysis after the cells were snap frozen. Our results could not be directly compared with the data in the previous study, also due to the difference in conditions; specifically, our assays were conducted at 25°C and 37°C, whereas the previous work was performed at standard laboratory conditions of 30°C.The impact on both transcription and translation was assessed through comparison of RNA-seq and Ribo-seq data by measuring translation efficiency with the deltaTE tool (Chothani *et al*, 2019). This allowed us to identify genes whose expression is altered primarily at the transcript level (differentially transcribed gene or DTG) and genes that show a change at the translation level (differential translation efficiency gene or DTEG) (Figure S5B-C). The deltaTE tool also allowed us to classify genes in four different categories: (1) translationally forwarded genes that exhibit a significant change in RNA-seq and Ribo-seq at the same rate, resulting in no impact on translation; (2) translationally exclusive genes that show a significant change in Ribo-seq, with no change in RNA-seq, impacting translation; (3) translationally buffered genes that exhibit a significant change in Ribo-seq that counteracts the change in RNA-seq, buffering the effect of transcription; and (4) translationally intensified genes that show a significant change in Ribo-seq that amplifies the effect of transcription in the same direction.

**Figure 5.**
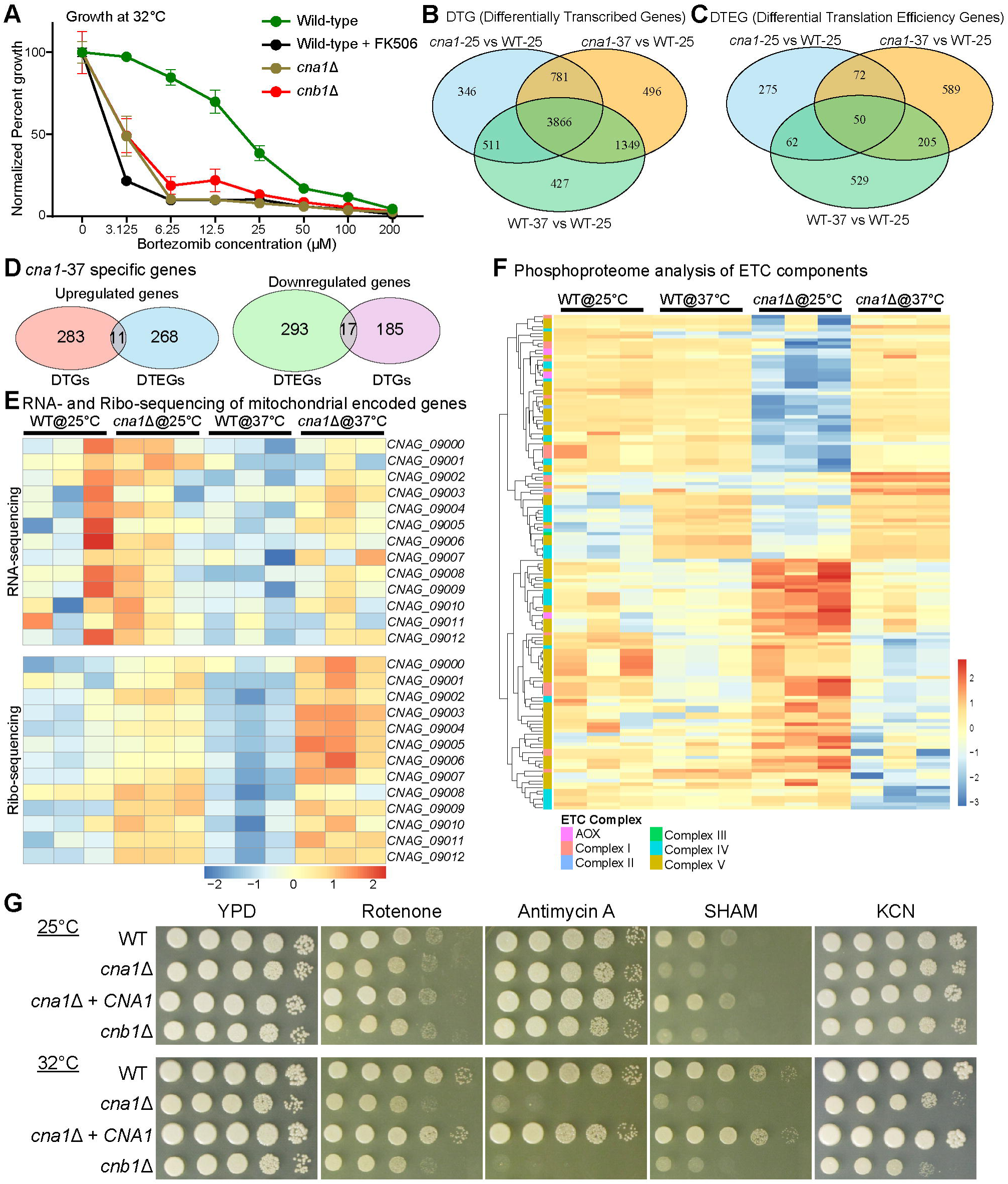
Calcineurin regulates mitochondrial function during thermal stress. **(A)** Growth pattern of various strains in the presence of proteasome inhibitor, Bortezomib. **(B-C)** Venn diagrams showing the overlap of genes that are differentially regulated at the transcriptional level (DTGs in A) or both the transcriptional and translational levels (DTEGs in B). **(D)** Venn diagrams showing the relationship between genes that are either up-or down-regulated in the *cna1*Δ mutant at 37°C exclusively. **(E)** A heatmap revealing the impact of calcineurin and temperature on the transcription and translation of mitochondrially encoded genes. **(F)** A heat map showing the calcineurin-dependent phosphopeptides at 37°C for proteins that are part of the electron transport chain (ETC) complexes. **(G)** Plate images showing the growth pattern of calcineurin mutants at two different temperatures in response to ETC complex inhibitors.

A pair-wise comparison of wild-type cells subjected to 37°C heat stress for 2 hours (WT-37), the *cna1*Δ mutant grown at 25°C (*cna1*-25), or the *cna1*Δ mutant with 2 hours of 37°C heat stress (*cna1*-37) with the wild-type cells grown at 25°C (WT-25) identified genes belonging to each of these categories for each condition (Figure S4B). To specifically define genes that are affected in the calcineurin mutant, we identified DTGs and DTEGs for each condition as compared to the control WT-25 group (Figure 5B-C). Further overlapping comparisons of three experimental groups (WT-37, *cna1*-25, and *cna1*-37) identified many DTGs and DTEGs in each category. Comparison of DTGs and DTEGs specific to the *cna1*-37 group identified genes that are up-or down-regulated (Figure ,5D, and Table S4). This analysis also revealed little overlap between DTGs and DTEGs, suggesting that calcineurin-mediated regulation of transcription and translation is relatively mutually exclusive.

### Calcineurin regulates mitochondrial function at two levels

Among the genes that exhibited significant changes in their translation in the *cna1*-37 group, we observed a cluster of mitochondrial-encoded genes (Figure 5E). Further analysis of all of the mitochondrial-encoded genes revealed an impact on their translation in the calcineurin mutant. This impact was observed in the calcineurin mutant grown at 25°C, but was significantly higher when grown at 37°C. Combined with the interactions of calcineurin with several mitochondrial proteins involved in energy metabolism detected by TurboID, we hypothesized that calcineurin regulates the function of the electron transport chain (ETC). Next, we analyzed the phosphoproteome data to identify if any of the ETC components are candidate substrates of calcineurin. Interestingly, several phosphopeptides belonging to the ETC complex proteins were found to be differentially phosphorylated in the calcineurin mutants (Figure 5F).

To further validate the involvement of calcineurin in mitochondrial function, we employed inhibitors of the ETC complexes and found that calcineurin mutants were hypersensitive to all of the ETC complex inhibitors in a temperature-dependent manner, albeit to different degrees (Figure 5G). The most severe sensitivity was observed with Antimycin A and SHAM, inhibitors of complex III and alternative oxidase, respectively. These results show that calcineurin plays an important role in mitochondrial function in this organism. We attempted to co-localize calcineurin with mitochondria by staining with MitoTracker in a GFP-Cna1 expressing strain, but were unable to confidently co-localize the two with or without heat stress. We also determined whether calcineurin might be acting on mitochondrial proteins during their import by dephosphorylating their cytosolic-exposed tails. Mapping of identified phosphopeptides along the protein lengths for ETC components did not indicate a specific pattern, and phosphopeptides were mapped throughout the protein lengths (Figure S6). This suggests that calcineurin’s role in this process may be an indirect one and occur at two different levels. First, calcineurin may dephosphorylate several of the ETC proteins, most likely prior to their import into the mitochondria. Second, calcineurin may regulate the translation of mitochondrial-encoded genes, several of which are key components of the ETC, and thereby impact mitochondrial function.

### Calcineurin activity is regulated at multiple levels through domains of its catalytic subunit

After defining the functional networks for calcineurin through genetic and physical interaction mapping, we next studied the structural regulation of calcineurin function. The calcineurin catalytic A subunit Cna1 in *C. neoformans* consists of several structural domains, including a long, highly disordered C-terminal tail and an N-terminal region preceding the predicted phosphatase domain (Figure 6A-B). Both of these regions are disordered and not resolved in the known *C. neoformans* calcineurin complex structure (Juvvadi *et al*, 2019), and neither is present in human calcineurin A. AlphaFold3 prediction of the calcineurin heterodimer (Cna1 and Cnb1) in complex with calmodulin (CaM), revealed a structure in accord with the known structures, as well as the interaction of Cnb1 and CaM with their respective binding regions on Cna1.

**Figure 6.**
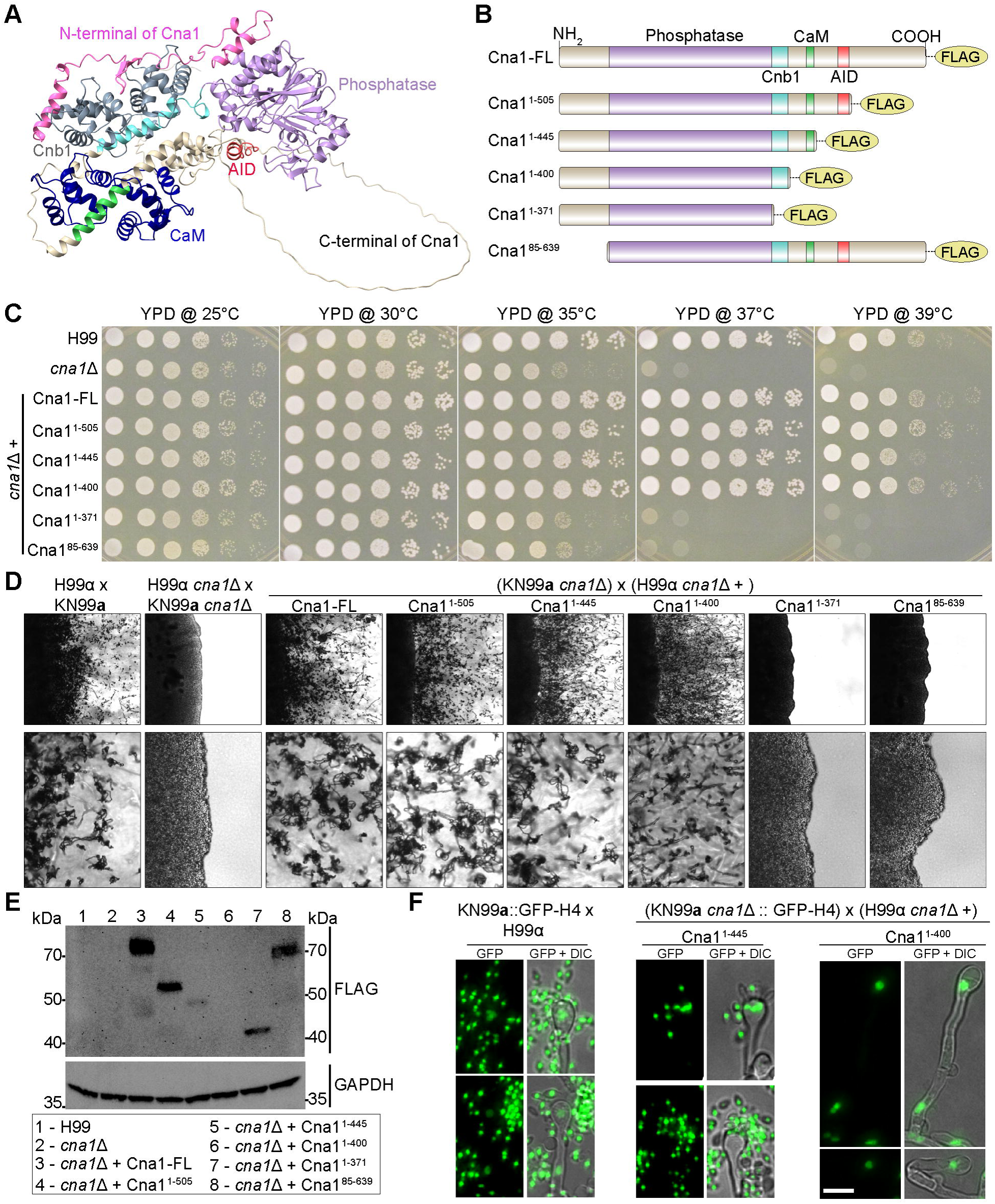
Calcineurin activity is regulated at multiple levels through domains of its catalytic subunit. **(A)** An AlphaFold prediction of the functional calcineurin complex consisting of the catalytic subunit Cna1, the regulatory subunit (Cnb1; Gray), and calmodulin (CaM; Blue). The specific domains within Cna1 are marked with different colors as follows: N-terminal (pink), phosphatase domain (purple), Cnb1-binding domain (cyan), CaM binding domain (green), and autoinhibitory (AID) domain (red). **(B)** A cartoon representation of Cna1 structural domains and truncation alleles generated. Cna1-FL refers to the full-length (FL) complementation allele, whereas the exact length for the rest of the alleles is marked with their retained amino-acid numbers. Cnb1 refers to the Cnb1 binding domain, CaM to the calmodulin-binding domain, and AID to the autoinhibitory domain. **(C)** Plate images of serial dilution spotting assays showing the growth complementation by each truncation allele compared to the growth of the calcineurin mutant and wildtype (H99) at various temperatures. **(D)** Microscopy images showing the functional complementation of sexual reproduction by Cna1 alleles. The upper panel presents a 4X magnification of mating assay spots after 2 weeks, whereas the lower panel depicts the 20X magnification of the same. **(E)** Western blotting showing the protein levels of calcineurin truncation alleles post-exposure to 37°C for 2 hours. **(F)** Fluorescence images showing a meiosis-specific defect of the Cna1^1-400^ allele, resulting in defective sporulation. Scale bar, 5 µm.

To test the roles of these domains in calcineurin function, alleles of calcineurin lacking one or more of the structural domains were generated and tested for function (Figure 6). This analysis showed that the disordered C-terminal region plays a minimal role during high-temperature growth (compare row 4 to rows 1 and 3 in Figure 6C). The strain expressing an allele lacking the autoinhibitory domain exhibited a growth defect at 39°C, suggesting that constitutively active calcineurin is detrimental to cells at high temperature (compare row 5 with row 3). Interestingly, this growth defect was mitigated upon further loss of the calmodulin-binding domain (compare row 6 with row 5), highlighting an important regulatory role through CaM binding. This finding also indicates that calmodulin binding is not essential for calcineurin’s function in the absence of the AID, similar to what has been previously observed in human cells and *Aspergillus fumigatus* (Juvvadi *et al*, 2013; Li *et al*, 2016). Surprisingly, the allele lacking the N-terminal region completely failed to complement growth at high temperature, similar to the Cnb1 binding domain allele (rows 7 and 8).

Western blot analysis of strains expressing these truncated derivatives revealed that the C-terminal domain plays a role in the stability of Cna1 at 37°C, and further removal of the autoinhibitory domain (AID) renders Cna1 highly unstable (Figure 6E). Additionally, abrogation of the CaM binding domain rendered truncated Cna1 undetectable in our experiments, but a protein with a further deletion of the Cnb1 binding domain was detected. The deletion of the N-terminal domain did not impact protein stability (Figure 6E).

Calcineurin was previously shown to localize to P-bodies/stress granules at 37°C in *C. neoformans*, and this may contribute to its function during thermotolerance (Kozubowski *et al*., 2011a). To study whether deletion of the structural domains impacts localization, additional strains were generated that express N-terminal mRuby3-tagged alleles in the same configuration from the safe haven locus, and calcineurin localization and function were assessed (Figure S7). Interestingly, the N-terminal mRuby3 tagging of these alleles led to differential complementation of Cna1 functions as compared to the C-terminal FLAG tag. Specifically, the strains expressing the C-terminal deletion allele showed relatively poor growth at higher temperature (Figure S7B). Co-localization analysis of these alleles in combination with a P-body marker (GFP-Dcp1) revealed that deletion of either the C-terminal or N-terminal domains partially impacted Cna1 localization to these structures, but neither was essential for its localization (Figure S7D).

### Calcineurin plays a calmodulin-dependent role during meiosis

Calcineurin plays an essential bilateral role in sexual reproduction of *C. neoformans* (Cruz *et al*., 2001). We determined the impact of domain truncations on sexual reproduction by crossing each of the truncation allele-expressing strains with a *cna1*Δ mutant of the opposite mating type. Similar to the thermotolerance phenotype, the allele lacking the C-terminal domain did not have an impact on calcineurin’s role during sexual reproduction, whereas the allele lacking the N-terminal domain did not restore any level of function (Figure 6D). Interestingly, the constitutively active calcineurin (column 5) did not have a substantial impact on mating, indicating differential regulation of calcineurin in sexual reproduction and 37°C growth. However, the deletion of the calmodulin-binding domain resulted in defective sporulation but normal hyphal growth, suggesting that calcineurin’s role in sporulation is calmodulin-dependent. A defect in sporulation could occur for multiple reasons, including a defect in nuclear fusion, meiosis, or the formation of spores itself. We tested this by crossing the strain expressing the Cna1 allele lacking the CaM binding domain with a GFP-H4 expressing *cna1*Δ mutant strain. Imaging of basidia in these crosses revealed that cells failed to undergo successful meiosis after nuclear fusion and harbor a single nucleus (Figure 6F). Basidia in crosses involving the CaM-binding domain-containing allele harbored four nuclei in a pre-sporulation or post-sporulation stage, similar to the cross of wild-type strains expressing GFP-H4. Combined, these results show that calcineurin functions in two steps during sexual reproduction in *C. neoformans*, first in hyphal elongation, which is calmodulin-independent, and second in meiosis, which is calmodulin-dependent.

Similar assays with the N-terminally tagged alleles revealed a striking difference for the allele lacking both the C-terminal and AID domains, which almost completely failed to complement the role of Cna1 during sexual reproduction, whereas the same allele without the N-terminal tag supported development similar to the wild-type (Figure 6D and S7C). Interestingly, the filamentation defect was restored to normal levels on further deletion of the CaM binding domain, but without sporulation, similar to the allele without the N-terminal tag. These results suggest that the N-terminal tag for the constitutively active Cna1 is deleterious for its function during sexual reproduction and further highlight the importance of the N-terminal region. Taken together, this analysis suggests that calcineurin activity is regulated at multiple levels through its structural domains. Furthermore, each of calcineurin’s roles is differentially regulated through these domains, and they contribute at different levels to multiple cellular functions.

### The C-terminal region of Cna1 plays an important role in virulence

Next, we determined the impact of these structural domains on the *in vivo* roles of calcineurin in a murine inhalation infection model. As expected, strains lacking calcineurin were completely avirulent, and function was restored to the wild-type level by expression of the full-length Cna1 allele (Figure 7A). Interestingly, strains expressing the allele lacking the C-terminal domain showed a delay in virulence, which was not further impacted by the removal of the AID and CaM-binding domains. As expected, the deletion of the Cnb1 domain or the N-terminal domain led to a complete loss of function, and these strains were avirulent. These results suggest that the disordered C-terminal region plays a specific role during infection that is separate from thermotolerance. Furthermore, the mild growth defects observed for strains expressing alleles lacking the AID or CaM binding domains in our thermotolerance assays may not be physiologically relevant, as their deletion did not impact virulence. Disordered regions have been proposed to play a role in CO_2_ sensing and response, a function that could explain the delay in virulence for the C-terminal lacking mutants. However, the strain lacking only the C-terminal region did not show an altered growth pattern in the presence of 5% CO_2_ in vitro as compared to the wild-type (Figure 7B). We also determined the impact of truncations on other virulence factors, such as melanin production (Figure 7C), and did not identify a defect in any of the truncation mutants.

**Figure 7.**
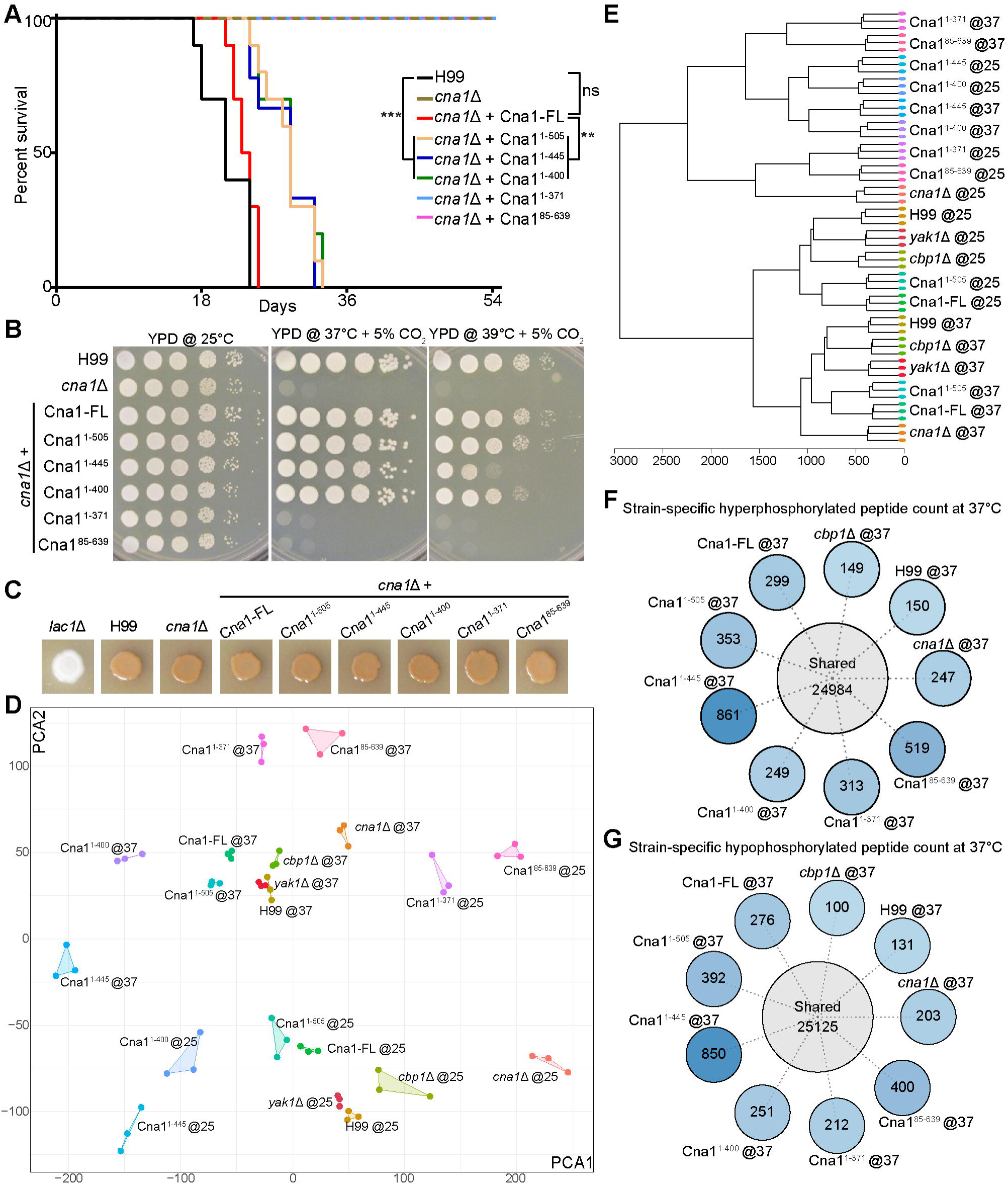
The C-terminal region of Cna1 plays a role in virulence. **(A)** A survival curve depicting the impact of Cna1 truncation alleles on virulence in mice. **(B)** Plate images showing growth complementation by truncation alleles in the presence of 5% CO_2_. **(C)** Images showing melanin accumulation in the strains expressing calcineurin A truncation alleles. **(D and E)** A PCA plot and dendrogram showing the clustering of phosphoproteome data from wild-type and calcineurin mutant strains at two different temperatures (25°C and 37°C). **(F and G)** Sunburst plots revealing the hyper-phosphorylated and hypo-phosphorylated peptide populations specific to each Cna1 truncation allele at 37°C.

Surprisingly, all of the strains expressing mRuby3-tagged Cna1 were found to be avirulent, including the strain expressing the full-length Cna1 allele, despite their growth at 37°C and normal production of virulence factors, such as melanin (Figure S8A-B). The C-terminal FLAG-tagged versions of the Cna1 alleles were expressed from the native locus, whereas the N-terminal mRuby3-tagged versions were expressed ectopically from a safe haven locus. We hypothesized that either the N-terminal tag, the expression locus, or both might be crucial for *in vivo* function. This was tested by integrating a tag-free version of full-length Cna1 either at the safe haven locus or next to the native locus. In both cases, virulence was largely restored, albeit modestly delayed at different levels compared to the wild-type (Figure S8C). These results strongly suggest that the N-terminal tag of Cna1 hinders *in vivo* function. Furthermore, the difference in virulence between strains expressing Cna1 from the native locus and the safe haven locus reveals that the genomic locus for Cna1 also plays a crucial role *in vivo* that is separate from its roles *in vitro*, and this finding warrants further investigation.

### Phosphoproteome analysis revealed domain-specific dephosphorylation substrates

To determine how the deletion of different domains impacts calcineurin activity, phosphoproteome analysis with strains expressing truncation alleles was conducted at 25°C and after 2 hours of heat stress induction at 37°C, along with the wild-type (H99) and calcineurin mutant (*cna1*Δ). In addition to the truncation alleles, we also included a previously characterized mutant lacking the calcineurin binding protein 1 (*cbp1*Δ) (Görlach *et al*, 2000). Cbp1 is a homolog of *Saccharomyces cerevisiae* Rcn1/calcipressin, which acts as a non-competitive calcineurin inhibitor (Hilioti & Cunningham, 2003). PCA analysis of these samples revealed a tight linkage between replicates, with a clear distribution of all 25°C samples in the lower right half and all 37°C samples in the upper left half of the plot (Figure 7D). The *cbp1*Δ and *yak1*Δ phosphoproteome was similar to the wild-type and suggests a minor role of Cbp1 and Yak1 in thermoregulation on their own, consistent with both previous findings as well as our results (Figure 1 and Fox & Heitman, 2005).

Further clustering analysis of all samples separated all truncation alleles except Cna1^1-505^ into a different cluster that harbored samples from both 25°C and 37°C (Figure 7E). Within this cluster, two alleles that showed the least separation based on temperatures were Cna1^1-445^ and Cna1^1-400^, both of which lack the AID and are functional. The alleles that were unable to complement functions (Cna1^1-371^ and Cna1^85-639^) were entirely separated based on temperature. Interestingly, while these two alleles appear identical to the *cna1*Δ mutant phenotypically, the phosphoproteome landscape in these strains was strikingly different. We hypothesize that this could be due to the presence of a truncated Cna1 protein that could interfere with some cellular processes. The two inactive truncated alleles include one that has the N-terminal region and the phosphatase domain, and the second, which lacks the N-terminal region but harbors all other domains. Because both of these forms are still expressed and can be detected by western blot and fluorescence microscopy (Figure 6E and S7D), they likely associate with other proteins or complexes and do so in a different manner than each other. Also, both of these truncation alleles express a form of calcineurin with the phosphatase domain that might act non-specifically on some proteins in the cell. As a result, these non-specific associations and dephosphorylation events for the truncated alleles lead to differences from the null allele.

Next, we identified allele-specific phosphopeptides that were enriched only at 37°C in each case (Figure 7F-G). Any peptide that was shared between any two samples, either at 25°C or 37°C, was classified as shared. As expected, the mutant lacking the AID (Cna1^1-445^) showed the maximum number of specific peptides. However, this was not limited to phosphopeptides that were downregulated in this mutant, but also included ones that were upregulated. We hypothesize that this might be due to a downstream effect, whereby the activity of other kinases and phosphatases is altered via constitutively active calcineurin. This might also explain the growth defect observed in this mutant at higher temperatures. Indeed, analysis of the specific peptides to identify unique proteins revealed the presence of several phosphatases and kinases that were differentially phosphorylated in this mutant (Figure S8D-E and Table S5).

## Discussion

The ability to grow at human body temperature is key to causing infections for environmental-dwelling pathogens such as *C. neoformans*. In this study, we examined the molecular mechanisms that govern thermotolerance in this critical priority fungal pathogen. Through a combination of genetic, biochemical, proteomic, and cell biology-based approaches, we defined a thermoregulatory network landscape in a human pathogen that functions uniquely in response to high temperature stress experienced during infection.

Isolation and analysis of genetic suppressors that bypass the requirement for calcineurin during high-temperature growth resulted in the identification of a majority of mutations in the kinase Yak1. Extension of this screen to a kinase deletion mutant library confirmed Yak1 and identified two additional kinases (Pan3 and Pka1), of which loss-of-function can partially bypass the requirement for calcineurin at high temperatures; however, Yak1 was found to be the primary kinase that acts antagonistically to calcineurin during growth at 37°C. Yak1 is a dual-specificity tyrosine-phosphorylation-regulated kinase (DYRK) that does not have a thermal stress phenotype in *C. neoformans* but shows reduced fitness during mouse infection (Boucher *et al*., 2025; Lee *et al*., 2016). Interestingly, a mutation in Yak1 was also identified in one (out of 64 isolates) isolate in a parallel genetic screen in the related species *Cryptococcus deneoformans* (Yadav *et al*., 2025).

DYRK-family kinases were also found to counter dephosphorylation of NFAT in a large genome-wide RNAi screen in *Drosophila* (Gwack *et al*, 2006). NFAT is the ortholog of the Crz1 transcription factor, which we identified as a shared substrate for both calcineurin and Yak1, suggesting that this calcineurin-Yak1 relationship might be evolutionarily conserved beyond *Cryptococcus*. In addition to Crz1 as a shared substrate, we also identified several other proteins involved in cell trafficking, microtubule dynamics, as well as several P-body proteins. This analysis suggests a broad antagonistic relationship between calcineurin and Yak1, which might have been missed in the study in *Drosophila* that focused on NFAT localization dynamics alone. Among other candidate kinases that suppress calcineurin function in our experiments, protein kinase A/Pka1 has also been shown to counter NFAT dephosphorylation by calcineurin, albeit indirectly via GSK3 (Sheridan *et al*, 2002). These studies suggest that calcineurin and kinase modules may be conserved across evolution, with conserved downstream effectors such as NFAT/Crz1 as well as additional effectors that may be species-specific.

When comparing the two sister species *C. neoformans* and *C. deneoformans,* Yak1 wasa found to be the only shared protein, mutations in which resulted in calcineurin bypass. *C. deneoformans* suppressors primarily harbored either dominant mutations in the genes encoding Imp2, Cts1, or Chs6 or segmental aneuploidy resulting in loss of Cdc42 (Yadav *et al*., 2025). The only similarity among these proteins is that Imp2 is dephosphorylated by calcineurin in both species and is one of the substrates shared by calcineurin and Yak1 (Figure 2E). Interestingly, the role for Cdc42 was found to be opposite in the two species, whereas loss of Cdc42 bypasses the requirement for calcineurin in *C. deneoformans* at 37°C; Cdc42 loss in *C. neoformans* results in hypersensitivity to calcineurin inhibition even at 25°C. These findings suggest that calcineurin signaling has been significantly rewired between these two species at the functional level. On the contrary, comparison of the calcineurin physical interactome as well as analysis of its substrates among these two species reveals a highly similar signaling network. We think that while the underlying signaling network mediated by calcineurin has remained conserved, its function has been specialized for cytokinesis in *C. deneoformans* but remains multi-faceted in *C. neoformans* (Mitchell, 2025; Yadav *et al*., 2025). How such a specialized function for a signaling cascade has been achieved in *C. deneoformans* remains an interesting question. A recent study in an evolutionarily distant fungal pathogen, *Aspergillus fumigatus,* identified that loss of function mutations in calcium channel subunits bypass calcineurin inhibition to allow for improved filamentous growth, which is regulated by calcineurin signaling (Lehmann *et al*, 2026). This suggests a broad conservation of mechanisms that bypass calcineurin in fungi and could be tested in other organisms. While this could be challenging due to the presence of several calcium channels in mammalian cells, suppression of the calcineurin signaling pathway was observed by L-type calcium channel blockers in rats (Liu *et al*, 2005), and similar approaches could assess the conservation of this association more broadly.

Phosphoproteome analysis identified several shared protein substrates between Yak1 and calcineurin that are involved in regulating microtubule dynamics, transcription, and protein modification, suggesting a co-regulation of several cellular processes between the two enzymes. Among these, microtubule components α-tubulin and β-tubulin were also identified as interacting partners of calcineurin with TurboID, indicating a potential role of calcineurin in microtubule dynamics. By fluorescence microscopy, we found a differential microtubule localization pattern for calcineurin mutants, with longer and more stable microtubule structures observed even at 25°C. Furthermore, calcineurin mutants were found to be hypersensitive to the microtubule-depolymerizing drug benomyl and exhibited unstable GFP-tubulin signals at 37°C. Calcineurin has been shown in neuronal cells to dephosphorylate tubulin and microtubule-associated proteins such as tau and MAP-2 and thus regulate microtubule assembly (Descazeaud *et al*, 2012; Goto *et al*, 1985). Recently, calcineurin was also shown in human cells to associate with centrosomes and regulate cilia length, which was originally detected through BioID proximity labeling assay (Tsekitsidou *et al*, 2023; Wigington *et al*, 2020). Our study further strengthens this link between calcineurin and microtubule dynamics and reveals a direct impact on genome stability and nuclear segregation.

With proximity ligation, we identified a strong association of calcineurin with the translation initiation machinery, including seven subunits of the eIF3 complex and two of three subunits of the eIF2 complex (Brito Querido *et al*, 2024; Brito Querido *et al*, 2020; Petrychenko *et al*, 2025). Furthermore, several translation initiation factors (TIFs) were also found to be candidate substrates for calcineurin, some of which were shared with Yak1 (Figure 2E). These findings corroborate our previous studies, which also showed some TIFs as candidate substrates for calcineurin (Park *et al*., 2016) and highlight a role for calcineurin in translation initiation and protein synthesis. Ribo-seq analysis in parallel with RNA-seq revealed that calcineurin indeed plays a major role in protein translation, in addition to its role in transcriptional regulation. GO analysis identified an impact on several cellular processes, including protein translation, cell cycle progression, and extracellular vesicle production. We note that our Ribo-seq dataset is not comparable to previously published studies in terms of quality and thus, we are unable to thoroughly characterize the impact of calcineurin on translation machinery and identify key genes whose translation is impacted. Our data, however, strongly suggests a key role for calcineurin in this process that needs to be investigated. Surprisingly, translation of the entire suite of genes encoded by the mitochondrial genome was found to be significantly decreased in the calcineurin mutant, suggesting a direct impact on mitochondrial function. Interestingly, our TurboID data also showed Cna1 interactions with several mitochondrial ETC proteins, many of which were found to be differentially phosphorylated in a calcineurin-dependent manner in the phosphoproteome data. Our attempts to co-localize calcineurin with mitochondrial markers, such as Mito-tracker, failed, and analysis of calcineurin peptide substrate location across protein lengths did not reveal a specific pattern. These results suggest that calcineurin may regulate mitochondrial function via dephosphorylation of proteins prior to their import into the mitochondria, similar to what has been shown in mammalian cell lines and *Drosophila* (Cereghetti *et al*, 2008; Marchesan *et al*, 2024). Calcineurin was shown to localize to the mitochondrial membrane in mice (Miyata *et al*, 2021), and it is possible that calcineurin localizes to the mitochondrial membrane in *C. neoformans* as well and this requires further investigation.

Calcineurin signaling is a highly conserved signaling cascade and our study delineates its functional and physical networks in a basidiomycetous fungus. The two other organisms where this pathway has been extensively studied at the cellular level are an ascomycetous budding yeast, *Saccharomyces cerevisiae*, and human cell lines (Goldman *et al*, 2014; Ulengin-Talkish & Cyert, 2023; Wigington *et al*., 2020). A comparison of these networks across these organisms reveals several overlapping as well as unique functions. Two of the most conserved associations for calcineurin are cytoskeletal elements such as microtubules and with cell trafficking machinery, both of which are crucial for cell growth, shape, signaling, and motility. Another conserved association seems to be between calcineurin and the nuclear pore complex, albeit this association seems to be less prominent in both fungal species. Inversely, calcineurin plays an important role in protein translation in both *S. cerevisiae* and *C. neoformans*, but the evidence for this in humans is unclear. Combined, this suggests that calcineurin signaling evolution is significantly tied to certain key cellular functions across kingdoms. It is important to note that the physical interactions for calcineurin and their evolution can be significantly different from its functional genetic interactions between these species, adding another layer of complexity.

The catalytic Cna1 subunit consists of several domains, including two fungal-specific highly disordered N-terminal and C-terminal extensions as compared to their human counterpart. Through the generation of domain truncation alleles, we discovered that calcineurin plays a calmodulin-dependent role during meiosis, the N-terminal is essential for its function, and the C-terminal plays an important role during infection. The in vitro phenotypic assays did not identify an attributable function for the C-terminal specific role, indicating an *in vivo* specific function for this domain. The allele lacking the N-terminal region did not complement any known phenotypes and was comparable to the deletion mutant or allele lacking the Cnb1 binding domain, suggesting a complete loss of function. The absence of this region in mammalian calcineurin suggests a more fungal-specific essential function that could be exploited for drug development and warrants further studies.

Finally, our phosphoproteome analysis also revealed a significant impact of temperature in regulating the phosphoproteome landscape in *C. neoformans*. This presents another layer of regulation in addition to gene expression changes as well as impact on alternative transcription start sites already known in *C. neoformans* (Chow *et al*., 2017; Dang *et al*., 2024; Janbon *et al*, 2014). Gene ontology analysis of the phosphoproteins that were differentially phosphorylated at 37°C as compared to 25°C revealed that proteins involved in cytoplasmic translation, microtubule-based movement, mRNA and tRNA metabolism, ribosome biogenesis, vesicle transport, and chromatin reorganization were hyperphosphorylated, whereas proteins involved in cell organization or biogenesis were specifically hypophosphorylated at 37°C. This suggests a dramatic functional regulation of these proteins via phosphorylation during heat stress. Given that phosphorylation is a known post-translational modification that affects protein stability, structure, and function (Newcombe *et al*, 2022; Zhong *et al*, 2023), the impact of phosphorylation needs to be investigated in detail for thermotolerance as well as for other pathogenesis-related mechanisms. It is possible that some of these changes might also be due to changes at the proteome level itself, which requires further investigation and systemic analysis. We hypothesize that such investigations will result in a better understanding of not only the biology of these key pathogens but will also reveal molecular mechanisms governing pathogenesis and identify novel candidates for drug development.

## Acknowledgements

We are grateful to Professor Praveen R. Juvvadi, University of Arkansas for Medical Sciences, and Connor Larmore, Duke University School of Medicine, for critical reading of the manuscript. We thank Dr. Erik Soderblom, Greg Waitt, and Tricia Ho at the Duke University Proteomics and Metabolomics Core Facility for phosphoproteome as well as Cna1-TurboID data acquisition and initial analysis. We also thank Dr. Devi Swain Lenz and the Duke University Sequencing and Genomic Technologies (SGT) Core Facility for Illumina sequencing. We are thankful to Ann (Bin) Hai from CD Genomics, where Ribo-seq experiments and sequencing were performed. We also thank Professor John Panepinto, University at Buffalo, SUNY, and Professor Joel McManus, Carnegie Mellon University, for suggestions and advice on Ribo-sequencing analysis. This work was supported by NIH/NIAID R01 awards AI039115-28, AI050113-20, and AI172451-04 awarded to JH. JH is Co-Director and Fellow of the CIFAR program Fungal Kingdom: Threats & Opportunities. This work was funded in part by a developmental grant to VY from the Duke University Center for AIDS Research (CFAR), an NIH-funded program (5P30 AI064518).

## Materials and Methods

### Strains and media

*Cryptococcus neoformans* reference strains H99α and KN99**a** (Nielsen *et al*, 2003; Perfect *et al*, 1993) were utilized for experiments conducted in this study. The strains were cultured in YPD (1% yeast extract, 2% peptone, and 2% glucose) liquid or YPD agar media at 25°C, 30°C, 32°C, 35°C, or 37°C, depending on the experimental requirements. For calcineurin inhibition conditions, calcineurin inhibitors FK506 and CsA were added to the YPD agar media at a final concentration of 1 µg/ml and 50 µg/ml, respectively. Gene deletion mutants and ectopic tagged strains were generated via homologous recombination with CRISPR following the TRACE protocol (Fan & Lin, 2018) or biolistic transformation (Toffaletti *et al*, 2024), and transformants were selected on YPD agar media supplemented with either 100 µg/ml nourseothricin or 200 µg/ml G-418. The strains and primers employed in this study are presented in Table S6. Strain construction for FLAG-tagged, TurboID, and truncation allele-expressing strains is described separately below in specific sections.

### Dilution spotting assays

Strains for serial dilution spot assays were grown in 5 ml YPD media overnight, and the growth was quantified by measuring OD_600_. For each strain, 5 OD equivalent cells were harvested, washed with 1 ml of autoclaved water, and finally resuspended in 1 ml of autoclaved water. The cells were 10-fold serially diluted to obtain dilutions of 10^-1^, 10^-2^, 10^-3^, 10^-4^, and 10^-5^. For each dilution, 3 µl of the cell suspension was spotted on the YPD media or YPD media supplemented with various drugs: FK506 (1 µg/ml), CsA (50 µg/ml), benomyl (2.5 µg/ml or 5 µg/ml), rotenone (50 µg/ml), Antimycin A ( 2 µg/ml), SHAM (10 mM), and KCN (5 mM). The plates were incubated at the respective experimental temperature conditions for 2 to 3 days before imaging with a Nikon camera. The images captured were processed with Adobe Photoshop and Adobe Illustrator.

### Isolation of genetic suppressors

Spontaneous genetic mutations that suppress calcineurin’s role for during high-temperature growth were isolated in either a wild-type strain H99 or an H99 *cnb1*Δ deletion mutant. The strains were first streaked to obtain single colonies, and 15 such independent single colonies were inoculated in 5 ml of YPD media for each strain. An aliquot of 200 µl of the overnight-grown cultures at 25°C was plated on either YPD agar containing both FK506 and CsA for H99 or YPD agar media for the H99 *cnb1*Δ strain and incubated at 37°C. After 7 days of incubation, the plates were screened for the presence of resistant colonies, and 4 to 6 colonies from each plate were streaked on fresh plates and incubated again for 3 to 4 days to validate the resistant phenotype. Once validated, one colony per plate was used for further analysis as an independent genetic mutant for calcineurin suppression, and a total of 16 suppressors were isolated through this approach. These strains were used for DNA preparation and whole-genome sequencing analysis.

### Whole genome sequencing and variant calling

A total of 16 calcineurin suppressors, along with the H99 *cnb1*Δ mutant strain, were subjected to Illumina whole-genome sequencing. Each strain was grown overnight in 5 ml YPD liquid media. Cells were harvested by centrifugation at room temperature, and the cell pellet was frozen at -80°C. The cells were lyophilized, and DNA extraction was performed using the MasterPure yeast DNA purification kit (Biosearch Technologies). The isolated DNA was checked for quality via agarose gel electrophoresis and NanoDrop and was quantified using Qubit. The DNA isolated was then submitted to the Duke Sequencing and Genomic Technologies core facility for sequencing on the Illumina Novaseq platform.

The sequencing reads obtained were checked for quality and mapped to the *C. neoformans* H99 reference genome available on FungiDB (https://fungidb.org/fungidb/app) with Bowtie2, with default parameters and allowing every read to map only once. The BAM files containing mapped reads were converted to TDF using IGVtools and visualized in IGV for the genome coverage plots and to identify events of aneuploidy. In parallel, the BAM files were analyzed with Geneious Prime for variant identification in each case. Single Nucleotide Polymorphism (SNP) events that were covered with at least 90 reads and were present in 90% of covering reads were scored as variants and were considered for analysis. Among these, SNPs that were shared between the suppressor isolates and the *cnb1*Δ mutant were not considered for further analysis. This analysis resulted in the identification of only one SNP in the protein-coding regions per isolate for the majority of the isolates. For isolates that did not present any SNP in their protein-coding regions and did not harbor any aneuploidy events, SNPs present in the introns were analyzed and considered. The mutations identified are summarized in Figure 1.

### Kinase mutant genetic screen

All strains from the protein kinase deletion library, consisting of 129 kinase mutants, were revived on YPD plates (Lee *et al*., 2016). All strains were streaked on YPD+FK506+CsA-containing plates in replicates along with the wild-type H99. One set of plates was incubated at 25°C, and another set was incubated at 37°C for 4 days to screen for mutants that exhibit growth by suppressing calcineurin function. The mutants that showed better growth as compared to the wild-type were identified and subjected to a second round of screening. For the second round, the strains were grown overnight in YPD media, and their growth in the YPD+FK506+CsA media was determined using serial dilution spotting assays described above for better assessment of the suppression phenotype. Additionally, 35°C was used in addition to 25°C and 37°C to score for the growth suppression phenotype. This resulted in the identification of three mutants (*yak1*Δ, *pka1*Δ, and *pan3*Δ) that showed better growth on YPD+FK506+CsA media at 35°C and 37°C than the wild-type.

### Phosphorylation assays

To assess the phosphorylation levels for proteins of interest, the proteins were tagged with 1X FLAG at the C-terminus and expressed from the native locus. The tagging constructs were generated via overlap PCR and *C. neoformans* was transformed with the TRACE method (Fan & Lin, 2018). The correct transformants were confirmed by PCR analysis and western blot analysis. Confirmed strains for each protein and a wild-type control strain were utilized for phosphorylation measurement assays. Specifically, the strains were grown at 30°C overnight in 5 ml cultures. The next day, each strain was inoculated into two fresh 50 ml cultures with a starting OD_600_ of 0.2 and allowed to further grow until 0.8-1 OD was achieved. At this step, one set of cultures was treated with 2 µg/ml FK506. Then, all the cultures were shifted to grow at 37°C for one hour before the cells were harvested and immediately frozen. The frozen cell pellets were lyophilized, followed by two rounds of bead beating at 4°C to make a fine powder from the cell pellets. Next, 1.5 ml of RIPA buffer containing protease inhibitor cocktail (cOmplete™, Mini Protease Inhibitor Cocktail) and phosphatase inhibitor mix (PhosSTOP™) was added into each tube, and the samples were subjected to two additional rounds of bead beating. The cell lysate mix was centrifuged at 13,000 rpm for 15 minutes at 4°C to remove cell debris. Out of the total cell lysate, 1 ml was incubated with 20 µl of FLAG M2 magnetic beads for 2 hours at 4°C on a rotator. For the beta-tubulin immunoprecipitation assays, the cell lysate was first incubated with 10 µl of anti-beta-tubulin antibody (Proteintech) for one hour, followed by incubation with 20 µl of Protein G beads for one hour. Following the bead incubation for each sample, the beads were pelleted on a magnetic stand, and then washed twice with RIPA buffer containing protein inhibitor cocktail and PhosSTOP for 5 minutes each. The bound proteins were eluted in 100 µl of elution buffer by heating at 95°C for 10 minutes. 25 µl of eluted sample or total cell lysate was used to assess the protein phosphorylation of respective proteins with 12.5% SuperSep™ Phos-tag™ Precast gels (FUJIFILM), blotted onto a PVDF membrane, and probed with anti-FLAG or anti-beta-tubulin antibody.

### Proximity ligation (TurboID) assay

To generate strains for TurboID assays, the previously described TurboID sequence was amplified from the plasmid received from Addgene (Plasmid #126050) (Larochelle *et al*, 2019). The TurboID was optimized for expression in *C. neoformans* via insertion of two introns within the TurboID encoding region between 157-158 bp and 555-556 bp, respectively. The intron sequences were chosen from intron #3 and #1, respectively, from the Actin gene (CNAG_00483), and were inserted into the final construct via overlap PCR. For TurboID alone expression, the TurboID gene was expressed via the Actin promoter from the safe haven locus, whereas the Cna1-TurboID fusion protein (without any linker) expression was driven via calcineurin’s native promoter at the native site. Strains expressing either TurboID alone or Cna1-TurboID fusion were grown overnight in 50 ml of biotin-free YNB media. The following day, 500 ml of fresh biotin-free YNB media was inoculated from the overnight culture with an initial OD600 of 0.2, and cells were grown for 4 hours at 25°C. For FK506-treated samples, 1 µg/ml of FK506 was added to the media and grown for another hour. The selected samples for heat stress were then shifted to 37°C and and all cultures were grown for an additional one hour. Biotin (50 µM) was added to all samples and grown for 3 more hours, following which the cells were harvested and frozen for lyophilization.

The lyophilized cells were subjected to bead beating twice for 1 min each with 5 min interval. The lysed cells were resuspended in RIPA buffer with protease inhibitor, and bead-beating was repeated twice more to allow for complete resuspension of cells in the buffer. The lysate supernatant was purified after centrifugation at 13,000 rpm for 15 mins at 4°C. The collected supernatant was transferred to Amicon Ultra-4kDa Millipore filter tubes and centrifuged to concentrate the lysate and also to remove the excess biotin from the supernatant. The purified lysate was then quantified using the BCA protein quantification assay, and 3 mg of total protein was immunopulldown assays using Streptavidin Dynabeads. Specifically,40 µl of Dynabeads were added to each sample and incubated at 4°C overnight on a roller. Next, the beads were separated using the magnetic stand and washed twice with RIPA buffer containing 1 mM DTT, and 2%SDS solution in 50mM Tris-Cl, pH 7.5, each. The bound proteins were then eluted using 50 µl of elution buffer (25mM Tris-Cl, pH 7.5, 100mM NaCl, 2% SDS, 5mM biotin, 10mM DTT) by incubating the mixture at 90°C for 15 minutes. The elution step was repeated, and the eluted samples were pooled together. A fraction of the eluate (10 µl) was tested on SDS-PAGE, followed by silver staining, whereas the remaining sample was submitted for mass-spectrometry analysis at Duke University Proteomics and Metabolomics Core Facility.

A total of 18 samples (triplicates of each TID25, CT25, CTF25, TID37, CT37, CTF37 as described in Figure 2B) were used for mass-spectrometry analysis. Samples were spiked with undigested bovine casein at a total of either 120 or 240 fmol as an internal quality control standard. Next, samples were reduced with 10 mM dithiolthreitol for 30 min at 80°C, alkylated with 20 mM iodoacetamide for 30 min at room temperature, then supplemented with a final concentration of 1.2% phosphoric acid and 1024 μL of S-Trap (Protifi) binding buffer (90% MeOH/100mM TEAB). Proteins were trapped on the S-Trap microcartridge, digested using 20 ng/μL sequencing grade trypsin (Promega) for 1 hr at 47°C, and eluted using 50 mM TEAB, followed by 0.2% FA, and lastly using 50% ACN/0.2% FA. All samples were then lyophilized to dryness. Samples were resolubilized using 12 μL of 1% TFA/2% ACN with 25 fmol/μL yeast ADH.

Quantitative LC/MS/MS was performed on 3 μL using a nanoAcquity UPLC system (Waters Corp) coupled to a Thermo Orbitrap Fusion Lumos high resolution accurate mass tandem mass spectrometer (Thermo) equipped with a FAIMSPro device via a nanoelectrospray ionization source. Briefly, the sample was first trapped on a Symmetry C18 20 mm × 180 μm trapping column (5 μl/min at 99.9/0.1 v/v water/acetonitrile), after which the analytical separation was performed using a 1.8 μm Acquity HSS T3 C18 75 μm × 250 mm column (Waters Corp.) with a 90-min linear gradient of 5 to 30% acetonitrile with 0.1% formic acid at a flow rate of 400 nanoliters/minute (nL/min) with a column temperature of 55°C. Data collection on the Fusion Lumos mass spectrometer was performed for three different compensation voltages (-40v, -60v, -80v). Within each CV, a data-dependent acquisition (DDA) mode of acquisition with a r=120,000 (@ m/z 200) full MS scan from m/z 375 – 1500 with a target AGC value of 4e5 ions was performed. MS/MS scans were acquired in the ion trap in Rapid mode with a target AGC value of 1e4 and max fill time of 35 ms. The total cycle time for each CV was 0.66s, with total cycle times of 2 sec between like full MS scans. A 20s dynamic exclusion was employed to increase the depth of coverage. The total analysis cycle time for each injection was approximately 2 hours.

Following UPLC-MS/MS analyses, data were imported into Proteome Discoverer 2.5 (Thermo Scientific Inc.), and individual LC-MS data files were aligned based on the accurate mass and retention time of detected precursor ions (“features”) using the Minora Feature Detector algorithm in Proteome Discoverer. Relative peptide abundance was measured based on peak intensities of selected ion chromatograms of the aligned features across all runs. The MS/MS data was searched against the TrEMBL *C. neoformans* H99 database, a common contaminant/spiked protein database (bovine albumin, bovine casein, yeast ADH, etc.), and an equal number of reversed-sequence “decoys” for false discovery rate determination. Sequest, with Infernys enabled (v 2.5, Thermo PD), was utilized to produce fragment ion spectra and to perform the database searches. Database search parameters included fixed modification on Cys (carbamidomethyl) and a variable modification on Met (oxidation). Precursor mass tolerances were 2.0 ppm, and product ion mass tolerances were 0.8 Da, with full trypsin enzyme rules required. Peptide Validator and Protein FDR Validator nodes in Proteome Discoverer were used to annotate the data at a maximum 1% protein false discovery rate based on q-value calculations. Note that peptide homology was addressed using razor rules in which a peptide matched to multiple different proteins was exclusively assigned to the protein that has more identified peptides. Protein homology was addressed by grouping proteins that had the same set of peptides to account for their identification. A master protein within a group was assigned based on % coverage.

Raw intensity values were used for filtering out peptides that were not measured in at least 2 unique samples (50% of a single group). After that filter, any missing data missing values were imputed using the following rules; 1) if only a single signal was missing within the group of three, an average of the other two values was used or 2) if two out of three signals were missing within the group of three, a randomized intensity within the bottom 2% of the detectable signals was used. A normalization was then applied to the data by excluding the highest and lowest 10% of the signals and then making the average of the remaining signals the same across all samples. To summarize to the protein level, all peptides belonging to the same protein were summed into a single intensity (Table S7). These normalized protein level intensities were used for the remained of the analysis, which was done using the DEP2 pipeline (Feng *et al*, 2023).

### Phosphoproteome analysis

The wild-type strain H99, a calcineurin *cna1*Δ, *yak1*Δ, *cbp1*Δ, and 6 strains expressing FLAG-tagged Cna1-truncation alleles were grown in 20 ml liquid YPD for 24 hours at 25°C in triplicate. The culture was then split into two halves of 10 ml cultures, and one set was further incubated at 25°C for 2 hours, whereas the second set was subjected to heat stress at 37°C for 2 hours. Cells were harvested from all samples by centrifugation and immediately frozen at -80°C. All 60 samples were then used for phosphoproteome analysis at the Duke University Proteomics and Metabolomics Core Facility.

Samples were supplemented with 100 µl of 8 M urea in 50 mM ammonium bicarbonate before protein extraction using a bead beater (2 rounds at 10 sec). Following quantification by Bradford assay, 165 µg of protein from each sample was used for further analysis. Each sample was spiked with 1 or 2 pmol bovine casein as an internal quality control standard. Next, the samples were reduced for 30 min at 32°C, alkylated with 20 mM iodoacetamide for 30 min at room temperature, then supplemented with a final concentration of 1.2% phosphoric acid and 824 μl of S-Trap (Protifi) binding buffer (90% MeOH/100mM TEAB). Proteins were trapped on the S-Trap micro cartridge, digested using 20 ng/μl sequencing grade trypsin (Promega) for 1 hr at 47°C, and eluted using 50 mM TEAB, followed by 0.2% FA, and lastly using 50% ACN/0.2% FA. All samples were then lyophilized to dryness. The phosphopeptide samples were resuspended in 80% acetonitrile and 1% TFA prior to TiO2 enrichment. Each sample was subjected to complex TiOx enrichment using GL Biosciences TiO2 tips and manufacturer-recommended protocols. Eluted phosphopeptides were then subjected to C18 stage tip cleanup. All samples were frozen and lyophilized.

Quantitative LC/MS/MS was performed on 4 µl of each sample using an EvoSep One UPLC system coupled to a Thermo Orbitrap Astral high-resolution accurate mass tandem mass spectrometer. Briefly, each sample loaded in EvoTip was eluted onto a 1.5 μm EvoSep 150 µm ID x 15cm performance (EveoSep) column using the SPD30 gradient at 55°C. Data collection on the Orbitrap Astral mass spectrometer was performed in a data-independent acquisition (DIA) mode of acquisition with r=240,000 (@ m/z 200) full MS scan from m/z 380-1080 with a target AGC value of 4e5 ions. Fixed DIA windows of 5 m/z from m/z 380 to 1080 DIA MS/MS scans were acquired in the Astral with a target AGC value of 5e4 and a max fill time of 8 ms. HCD collision energy setting of 27% was used for all MS2 scans. The total analysis cycle time for each sample injection was approximately 40 min.

Following 61 total UPLC-MS/MS analyses (60 experimental samples and 1 internal quality control sample), data were imported into Spectronaut (Biognosis), and individual LC-MS data files were aligned based on the accurate mass and retention time of detected precursor and fragment ions. Relative peptide abundance was measured based on MS2 fragment ions of selected ion chromatograms of the aligned features across all runs. The MS/MS data were searched against a *C. neoformans* H99 database (downloaded in 2024), a common contaminant/spiked protein database (bovine albumin, bovine casein, yeast ADH, etc.), and an equal number of reversed sequence “decoys” for false discovery rate determination. A library was generated with only the LC-MS data files collected within this study. Database search parameters included fixed modification on Cys (carbamidomethyl) and variable modification on Met (oxidation), Protein N-term (acetyl), and Ser/Thr/Tyr (phos). Full trypsin enzyme rules were used along with 10ppm mass tolerances on precursor ions and 20ppm on product ions. Spectral annotation was set at a maximum 1% peptide false discovery rate based on q-value calculations. Note that peptide homology was addressed using razor rules in which a peptide matched to multiple different proteins was exclusively assigned to the protein that had the most identified peptides.

The output raw peptide intensity values from the Spectronaut detection software were noted. At this stage, any peptide that was not detected a minimum of 2 times across all of the samples and not detected in at least 50% of one of the biological groups was removed from further analysis. For remaining peptides, any missing data values were imputed using the following rules: 1) if less than 50% of signals were missing within a group, a randomized value near the average of the remaining values was calculated, or 2) if >50% of the signals were missing within the group, a randomized intensity within the bottom 1% of the detectable signals was used. Next, non-phosphorylated peptides and phosphopeptides that did not pass a 75% confidence of localization were removed from the analysis. The remaining phosphopeptides were then subjected to a robust mean normalization in which the highest and lowest 10% of the phosphopeptide signals were ignored, and the average value of the remaining phosphopeptides was used as the normalized factor across all samples. We then summed all of the same precursor states for the same phosphopeptide (i.e., different charge states) into a single modified phosphopeptide value and calculated the % coefficient of variation across each group. Following database searching and peptide scoring using a library-based Spectronaut search, the data was annotated at a 1% peptide false discovery rate. A total of 56,135 phosphopeptides were identified in the dataset corresponding to 2,992 phosphoproteins.

The normalized values (Table S8) were then used for analysis to calculate the fold-change between various groups, and significantly enriched peptides in each group were identified using DEP2 (4). Analysis was done using R scripts to identify the phosphopeptides that were significantly enriched in any specific group, depending on the analysis. All peptides that exhibited at least a 1.5-fold difference and a p-value < 0.05 were considered for the analysis.

### RNA-sequencing and alternative splicing

Three colonies for wild-type H99, *cna1*Δ, and *cnb1*Δ mutants were inoculated in 8 ml YPD medium and grown overnight at 25°C. The next day, each culture was split in half, and one set of these was incubated at 25°C, whereas the second set was incubated at 37°C for an additional 2 hours. Cells were then harvested, frozen, and subjected to lyophilization overnight. A total of 60-70 mg of lyophilized material was used for RNA isolation following the mirVana miRNA Isolation Kit manufacturer’s instructions for total RNA purification. RNA quality was determined using agarose gel electrophoresis, and it was quantified with a Qubit 3 Fluorometer before submitting for sequencing at the Duke University Sequencing and Genomic Technologies Core facility. RNA libraries were prepared with a KAPA HyperPrep RNA library kit, and 2 x 150 bp reads were sequenced on the Illumina NovaSeq X Plus.

Post sequencing, the reads were processed by trimming and checked for quality using trim_galore equipped with cutadapt (v 3.5). The trimmed reads were then mapped to the reference H99 genome STAR aligner in a two-step fashion; first, the reads were mapped to the rDNA as a reference, and reads not aligned to the rDNA sequence were extracted and then aligned to the H99 genome reference. The .bam files generated as a result were used for splicing variant analysis using rMATS-Turbo (Shen *et al*., 2014). The rMATS output files were analyzed using custom R scripts to identify condition-specific splicing events.

### Ribo-sequencing and translation efficiency analysis

Three independent colonies were grown overnight in the 50 ml YPD liquid media at 25°C for each of the wild-type H99 and the *cna1*Δ mutant. The next day, the culture was split into two halves, each with 25 OD equivalent of cells. One half was transferred to 37°C and further grown for 2 hours, and the other half was maintained at 25°C. Each culture was then split into two parts, one with an equivalent of 20 OD cells (for Ribo-seq) and another with 5 OD cells (for RNA-seq), before cell harvesting. The cell pellets were frozen in liquid nitrogen for 2 hours and then shipped to CD Genomics (https://www.cd-genomics.com/) using a dry ice shipment, where the following steps were performed.

For Ribo-seq samples, cells were fixed with harringtonine (final concentration of 2 μg/ml) at room temperature for 2 minutes, followed by treatment with cycloheximide (final concentration of 100 μg/ml) at room temperature for 1 minute. The treated cells were washed with 1X PBS, and 1 ml of lysis buffer was added and incubated on ice for 10 minutes. The cells were centrifuged at 17,000 g for 10 minutes at 4°C, and the supernatant was collected. The supernatant was then treated with 10 µl of RNase I and 6 µl DNase I at room temperature for 45 minutes. The samples were then used for ribosome separation using MicroSpin S-400 columns. The RNase-DNase treatment supernatant was added to the column and centrifuged at 600 g for 2 minutes at room temperature to collect the flow-through. Next, the ribosome-protected fragments (RPFs) were extracted by mixing 700 µl of lysis reagent with the flow-through and incubating at room temperature for 8 minutes. 140 µl of chloroform was added, mixed, incubated at room temperature for 5 minutes, and then centrifuged at 12,000 g for 15 minutes at 4°C. 350 µl of the supernatant was transferred to a new 1.5 ml tube, and 525 µl of ethanol was added and mixed. RPFs were purified using a column and finally eluted in 30 µl of RNase-free water.

For RNA-seq samples, the cells were resuspended in Trizol solution and RNA extraction was performed using the standard Trizol extraction protocol. The mRNA population was enriched using oligo-dT-mediated purification. The RNA extracted, as well as RPFs, were then checked for quality and used for 150-PE Illumina library preparation and sequenced using the Illumina HiSeq X Ten platform. 150-PE sequencing was employed for both RNA-seq and Ribo-seq, even though that is not necessary for Ribo-seq due to CD Genomics policy and to maintain the procedure uniformity. At the analysis step, only the READS1 set was used for Ribo-seq analysis and second pair was not analyzed. The reads obtained were processed by quality trimming with trim_galore equipped with cutadapt (v 3.5). The trimmed reads were then mapped to the reference H99 genome STAR aligner in a two-step fashion; first, the reads were mapped to the rDNA and U3 DNA sequence as a reference to filter out these reads. The remaining reads were aligned to the H99 genome reference to obtain the .bam files. The .bam files were used for ribo footprint measurement presented in Figure S5 using a custom R script. These .bam files were processed using the “bedtools coverage” tool to obtain read counts for each gene. These counts obtained for each sample were then combined for all RNA-seq samples and Ribo-seq samples separately to generate two gene count matrices with information for each gene and sample (Table S9). These two gene count matrices were then used for assessing impact on translation using the delta TE tool in R (Chothani *et al*., 2019). The PCA plots were also generated using these gene count matrices. The scripts used for this analysis are available on GitHub and can be accessed using the link provided in the Data availability section below.

### Generation of truncation allele strains

Calcineurin catalytic A subunit, Cna1, sequence was retrieved from FungiDB (CNAG_04796), and its various domains were mapped using Interpro and its alignment with the human calcineurin A sequence. Primers were designed to amplify allele-specific regions and placed under the native promoter consisting of 1000 bp upstream of Cna1 ATG. For FLAG-tagged strains, the C-terminal FLAG tag encoding sequence was inserted in the primer, which was then fused with a short terminator sequence and the selection marker G-418 sequence. The constructs were transformed into a *cna1*Δ mutant, and the promoter region and downstream regions were used for homologous recombination via the CRISPR-Cas9-mediated transformation method. For mRuby3-tagged alleles, the mRuby3 sequence was amplified from the donor plasmid containing the mRuby3 ORF and was inserted between the promoter and ATG. The final constructs were cloned into a plasmid designed for targeted insertion at the safe haven 2 locus in *C. neoformans* (Upadhya *et al*, 2017). Each plasmid harboring the truncation- and tagged alleles of Cna1 was then transformed into *C. neoformans* H99 as well as in the *cna1*Δ mutant using CRISPR-Cas9, and transformants with correct integration were identified using PCR analysis.

### Sexual reproduction

Mating assays to study the impact of Cna1 truncations on sexual reproduction were performed on MS media at room temperature. The parental strains of opposite mating types for each cross were grown on YPD agar plates for 2 days. An approximately equal number of cells for each parent was harvested from the plate and mixed together in 200 µl of distilled and autoclaved water. The cell suspension was mixed thoroughly by vortexing, and 3 µl of the cell suspension was spotted on the MS media plates and incubated for 2 weeks at room temperature. For fluorescence imaging assays, the plates were incubated for 3 weeks to allow for longer hyphal growth and separation from the yeast cells contained within the mating spot.

### Fluorescence microscopy

Cells expressing fluorescently labeled proteins were grown overnight in 5 ml of YPD liquid. The next day, 1 OD of cells was transferred to 5 ml of fresh YPD media and grown for 4 to 5 hours either at 25°C or 37°C, depending on the experimental condition. The cells were harvested and washed once with 1 ml of purified water. Finally, cells were resuspended in 200 µl of YNB media with 2% glucose and further incubated under the respective temperature conditions. From this, 5 µl of cell suspension was spotted onto a 2% agarose slab prepared in YNB+2% glucose media, which was then inverted into a glass-bottom MatTek dish. The cells were imaged with the DeltaVision microscope system at the Duke Light Microscope Core Facility equipped with an incubation chamber to allow for temperature-dependent imaging. The images were captured with mCherry and GFP filters for mCherry and GFP, respectively. For imaging of sexual hyphae and basidia, the mating patch was excised from the MS plate and was directly inverted on the MatTek dish for imaging. The images were then processed by SoftWoRx software, v 6.1, connected to the microscope and ImageJ. The processed images were assembled with Adobe Photoshop and Adobe Illustrator for presentation purposes.

### Mouse survival assays

Experiments utilizing mice were conducted in compliance with guidelines issued by the US Animal Welfare Act and by the Duke Institutional Animal Care and Use Committee (IACUC). All procedures involving animals were approved by Duke IACUC Duke’s Division of Laboratory Animal Resources (DLAR) under protocol no. A098-22-05. Wild-type *C. neoformans* H99, *cna1*Δ, and truncation allele expressing strains were grown in YPD medium overnight at 25°C in a roller drum. Cells were pelleted at 3,000 rpm and washed 3 times with sterile 1X PBS. Cells were counted for each strain with a hemacytometer and diluted to obtain 4 × 10^6^ cells/ml in PBS. An aliquot of 25 µl (equivalent to 10^5^ cells) was used for pulmonary inhalation infection of mice. Specifically, ten 3-4-week-old A/J mice (5 male and 5 female; Jackson Laboratory) were used for infection per strain. Mice were anesthetized utilizing an isoflurane chamber. While anesthetized, 25 μl of cell suspension was dropped onto the nares of mice, allowing them to inhale the full inoculum. Mice were observed for normal activity post-infection and monitored daily afterwards. All mice were weighed on Day three post-infection, which was used as the baseline weight for each mouse. Post 14 days of infection, mice were weighed daily and monitored for health and survival until reaching a humane endpoint, at which point they were sacrificed. The experiment was terminated at day 55 or 65, and any surviving mice were sacrificed. Survival analysis was plotted in GraphPad Prism, and statistical significance was determined using log-rank (Mantel-Cox) tests.

### Data availability

The whole genome sequencing data for all the suppressors and calcineurin mutant strains are available via NCBI BioProject PRJNA1306639. The mass spectrometry proteomics data have been deposited to the ProteomeXchange Consortium (Deutsch *et al*, 2023) via the PRIDE (Perez-Riverol *et al*, 2025) partner repository with the dataset identifiers PXD067478 (Cna1-TurboID data) and PXD069393 (Phosphoproteome data). All custom scripts used are available in github (https://github.com/vikasyadavsci/CN_thermoregulatory_network).

## Supplementary figure title and legends

**Figure S1. Genetic suppression analysis identified Yak1 as a major suppressor of calcineurin function.**

**(A)** A schematic detailing the protocol employed to isolate and analyze the spontaneous mutants that suppress the requirement for calcineurin function during high-temperature growth. **(B)** Plate images showing the growth of suppressors under different conditions as compared to the parent wild-type or *cnb1*Δ mutant strains. **(C)** Genome coverage heatmaps showing the aneuploidy events in four of the isolates that did not harbor any mutations in protein-coding regions. Each row represents sequencing data from one strain, with H99 *cnb1*Δ exhibiting uniform coverage throughout the genome as a control. The remaining four rows present suppressor isolates with at least one event of aneuploidy marked by a darker color depicting amplification of the region. **(D-E)** Plate images showing the growth of mutants of the calcium channel (D) and kinases identified through the screen of the kinase deletion collection (E) in the presence of the calcineurin inhibitors, FK506 and CsA. Two independent knockout strains for each kinase deletion mutant that are part of the kinase deletion collection are presented in panel E.

**Figure S2. Phosphorylation status of candidate substrates.** Western blot analysis showing the phosphorylated state of immunoprecipitated proteins in each case. The assays were performed with Phos-tag gels, which resolve the phosphorylated form of the protein distinctly, as observed for both the FLAG-tagged proteins. The treatment with FK506 is denoted by the presence of “+” and “-” in each case.

Figure S3. TurboID proximity ligation revealed novel calcineurin interactions.

**(A)** Western blot analysis showing the expression levels of TurboID-myc and Cna1-TurboID-myc at different temperatures. **(B)** Serial dilution spotting assays showing growth of TurboID-myc and Cna1-TurboID-myc expressing strains at different temperatures. **(C)** Images showing filamentation for various crosses. **(D)** Western blot analysis showing the level of biotinylation in different strains and conditions with or without 50 µg/ml biotin in the media. **(E)** A workflow depicting the development of the TurboID assay in *C. neoformans*.

**Figure S4. Calcineurin orchestrates several cellular processes.**

**(A)** A Venn diagram showing the overlap of Cna1-interacting proteins with *C. neoformans* proteins with LxVP or PxIxIT motifs. **(B)** Fluorescence images showing localization dynamics of GFP-tubulin in the wild type and calcineurin mutants (*cna1*Δ and *cnb1*Δ) at 25°C. Scale bar, 5 µm. **(C)** Fluorescence images showing the nuclear segregation defects in the calcineurin mutants at 37°C as compared to the wild-type. Scale bar, 5 µm. **(D)** A scheme showing the workflow employed to detect altered splicing events in calcineurin mutants. **(E)** An UpSet plot showing the relationship between representative splicing events in calcineurin mutants compared to wild-type cells.

**Figure S5. Ribo-sequencing analysis for calcineurin mutants.**

**(A)** Read length plots for mapped Ribo-seq datasets presenting the lengths for ribosome-protected fragments for each sample. Three replicates for each condition are presented here (marked as Rep 1, 2 and 3). **(B)** PCA plots showing the replicate correlation for both Ribo-sequencing and RNA-sequencing samples. The plots were generated after variance stabilizing transformation (VST) on the mapped sequencing read counts matrix generated for the DEseq2 analysis for both RNA-seq and Ribo-seq and plotted using PC1 and PC2 from the transformed matrix **(C)** Scatter plots showing the impact on transcription (RNA-seq) and translation (Ribo-seq) in the wild-type and calcineurin mutant during thermal stress, as compared to the wild-type at 25°C (WT-25).

**Figure S6: Phosphopeptide mapping for mitochondrial proteins.**

A map showing the phosphopeptide locations along the respective protein length for ETC components. Each protein was divided into 20 bins of equal length, and the position of each identified phosphopeptide was mapped.

**Figure S7.** Calcineurin activity is regulated at multiple levels through domains of its catalytic subunit.

**(A)** A cartoon representation of Cna1 structural domains and truncation alleles generated with an N-terminal mRuby3 tag. Cna1-FL refers to the full-length complementation allele, whereas the exact length for the rest of the alleles is marked with their amino-acid numbers. Cnb1 refers to the Cnb1 binding domain, CaM to the calmodulin-binding domain, and AID to the autoinhibitory domain. **(B)** Plate images of serial dilution spotting assays showing the growth complementation of each truncation allele compared to the calcineurin mutant at different temperatures. **(C)** Mating assays presenting the functional complementation of calcineurin function during sexual reproduction by truncation alleles with the N-terminal mRuby3 tag. The upper panel presents a 4X magnification of mating assay spots after 2 weeks, whereas the lower panel depicts the 20X magnification. **(D)** The localization pattern of mRuby3-tagged alleles of Cna1 after growth at 37°C for 30 minutes. GFP-Dcp1 served as a marker for the P-body. Bar, 5 µm.

**Figure S8.** The C-terminal region of Cna1 plays a role in virulence.

**(A, C)** Survival curves depicting the defective virulence potential of strains expressing calcineurin alleles harboring the N-terminal mRuby3 tag or when integrated at different locations in the genome. **(B)** Images showing melanin accumulation in strains expressing the Cna1 truncation alleles. **(D-E)** UpSet plots showing the relationship among proteins that are differentially hyper-phosphorylated (D) and hypo-phosphorylated (E) in strains expressing the Cna1 truncation allele series.

## Supplementary Information

**Table S1.** List of shared proteins that are hyperphosphorylated in *cna1*Δ mutant and hypophosphorylated in *yak1*Δ mutant.

**Table S2.** List of Cna1-TurboID interacting proteins.

**Table S3.** List of alternatively spliced genes in calcineurin mutants.

**Table S4.** List of genes that are differentially regulated at the transcription and translation level in the *cna1*Δ mutant at 37°C.

**Table S5.** List of proteins hyper- and hypo-phosphorylated in Cna1 truncation allele mutants.

**Table S6.** Strains and primers used in this study.

**Table S7.** Normalized quantification data for TurboID assay samples.

**Table S8.** Normalized quantification data for phosphoproteome samples.

**Table S9.** Data tables for RNA-seq and Ribo-seq reads mapping statistics and DEseq2 analysis.

