## Supplementary figures and images for "Thermoregulation network governing virulence of a critical human fungal pathogen"

### Supplementary Figure S1

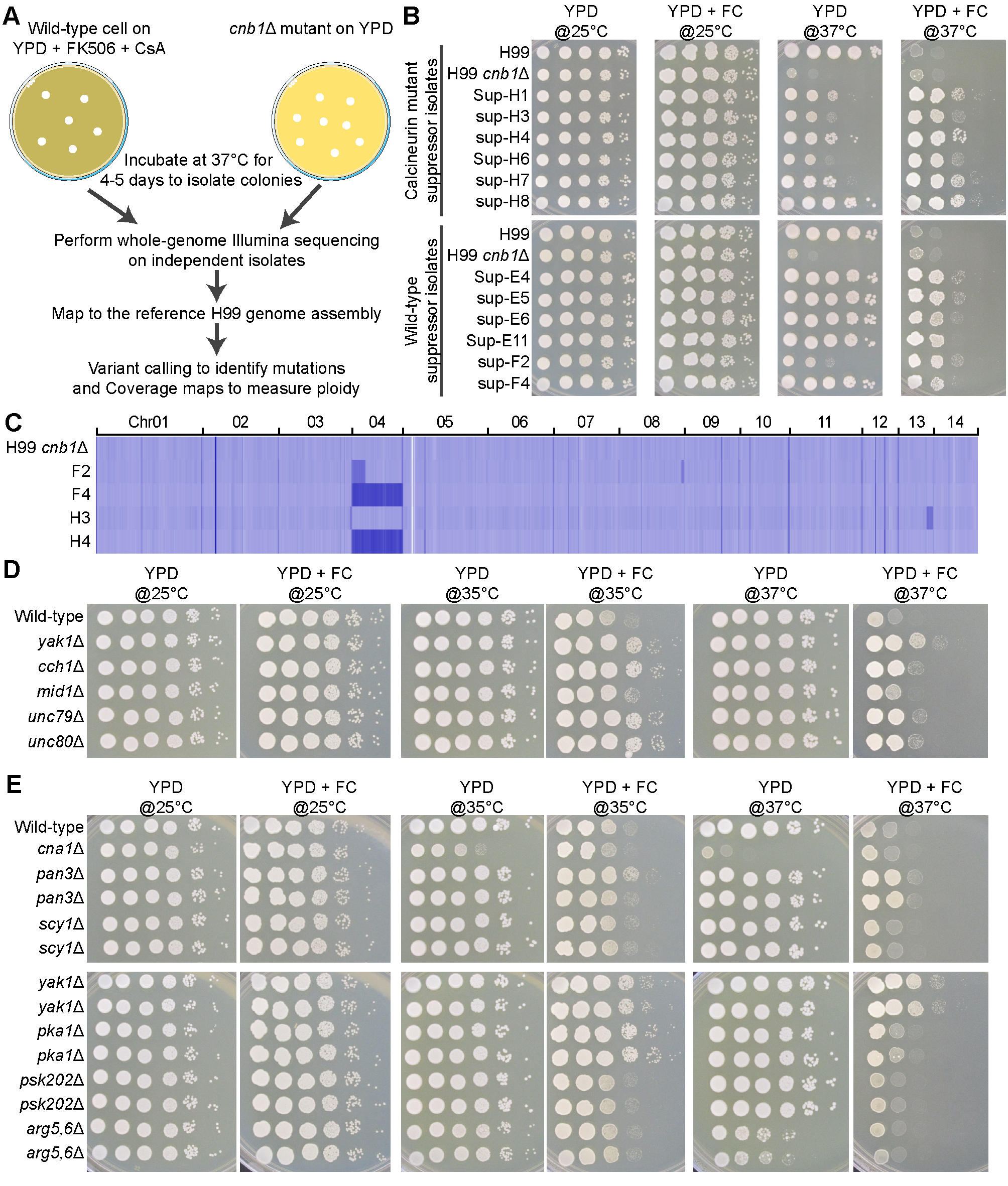

### Supplementary Figure S2

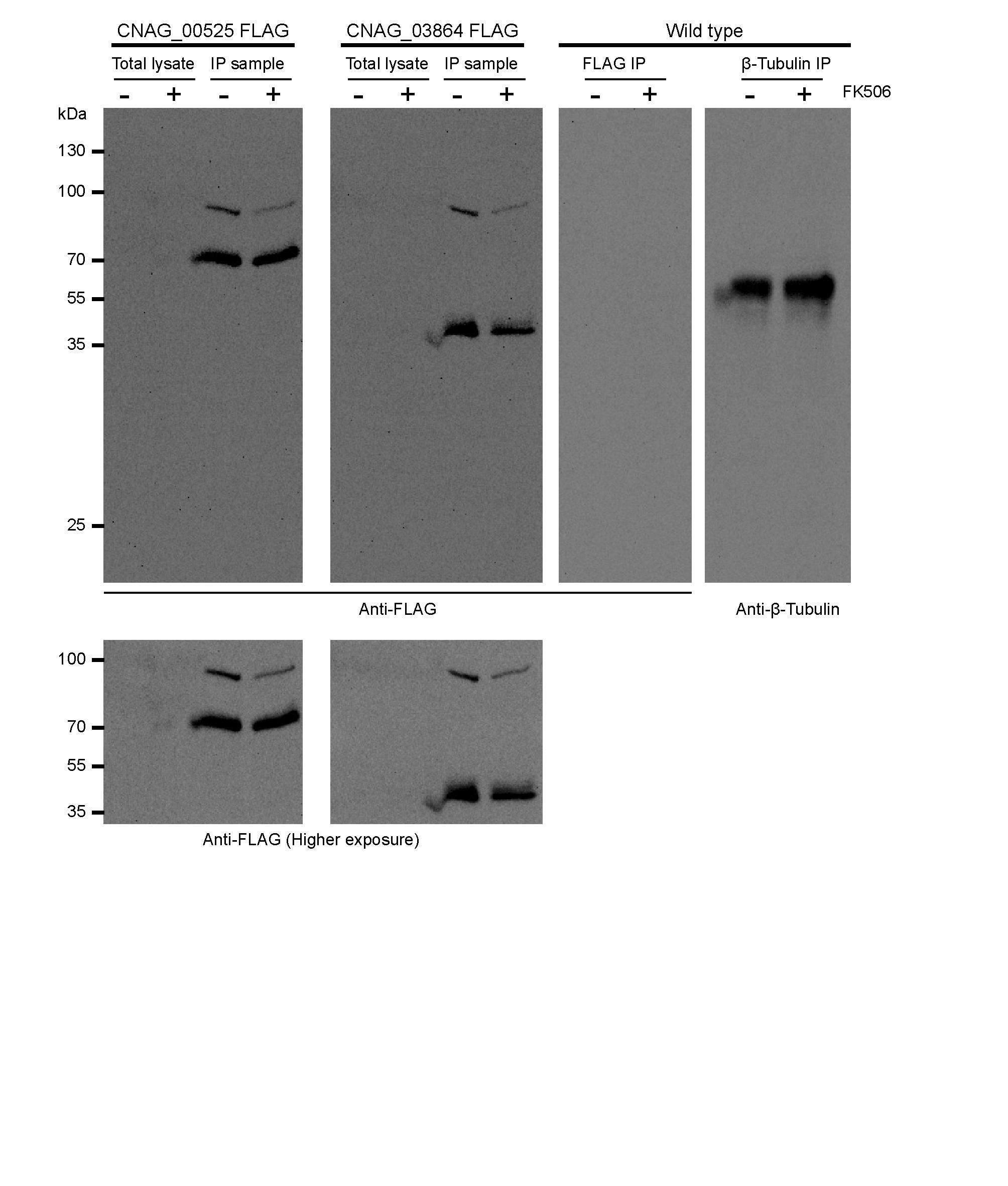

### Supplementary Figure S3

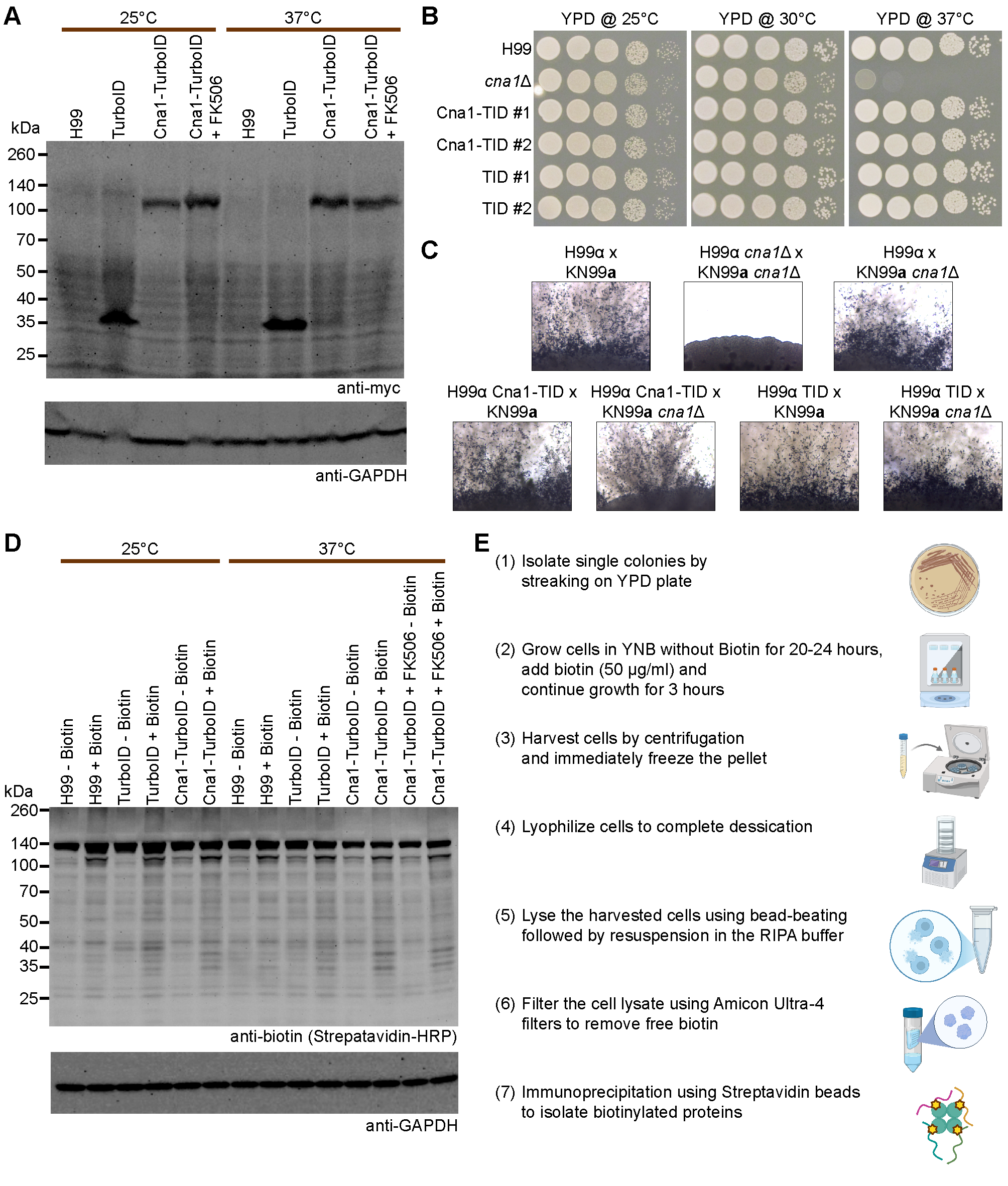

### Supplementary Figure S4

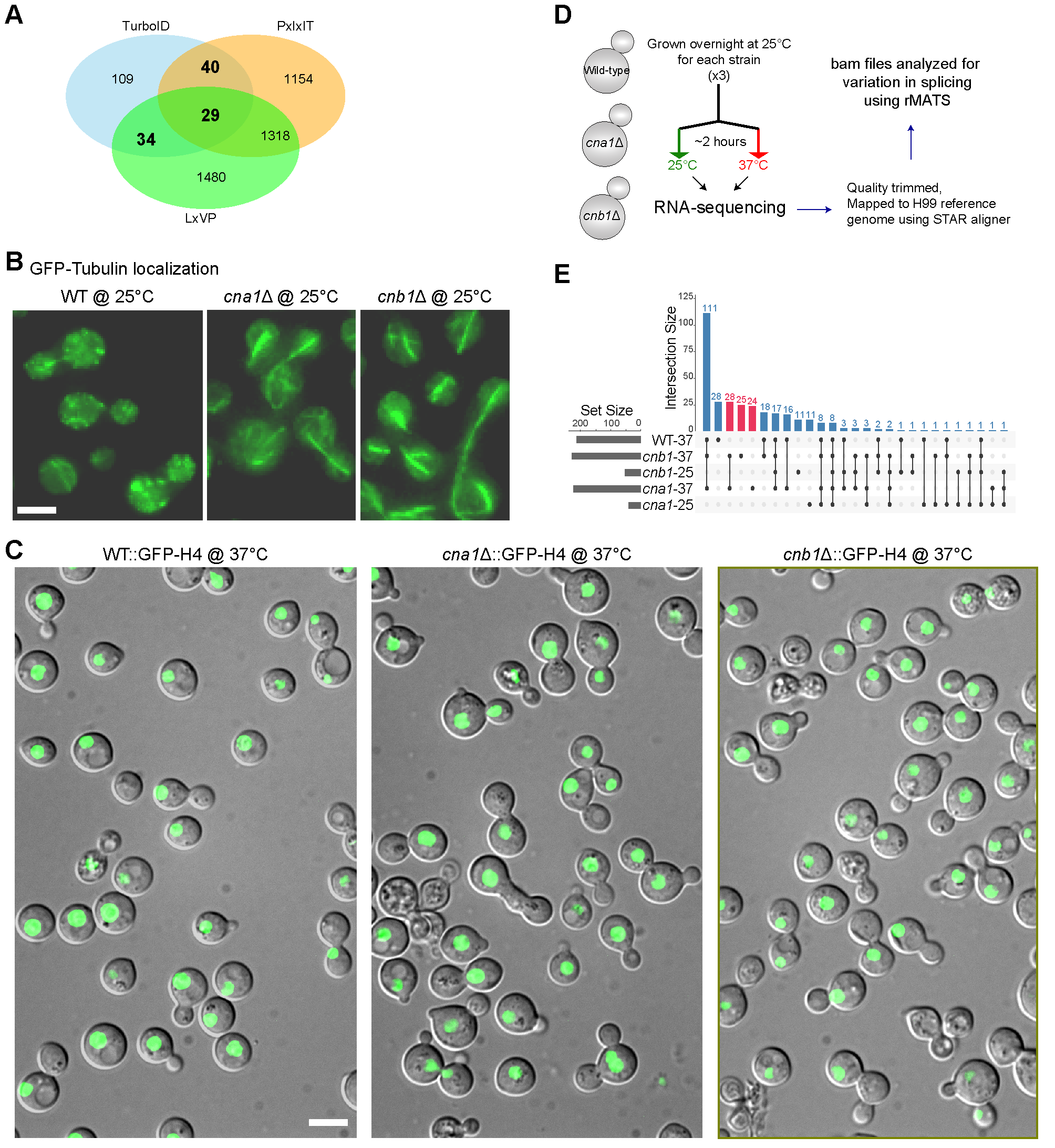

### Supplementary Figure S5

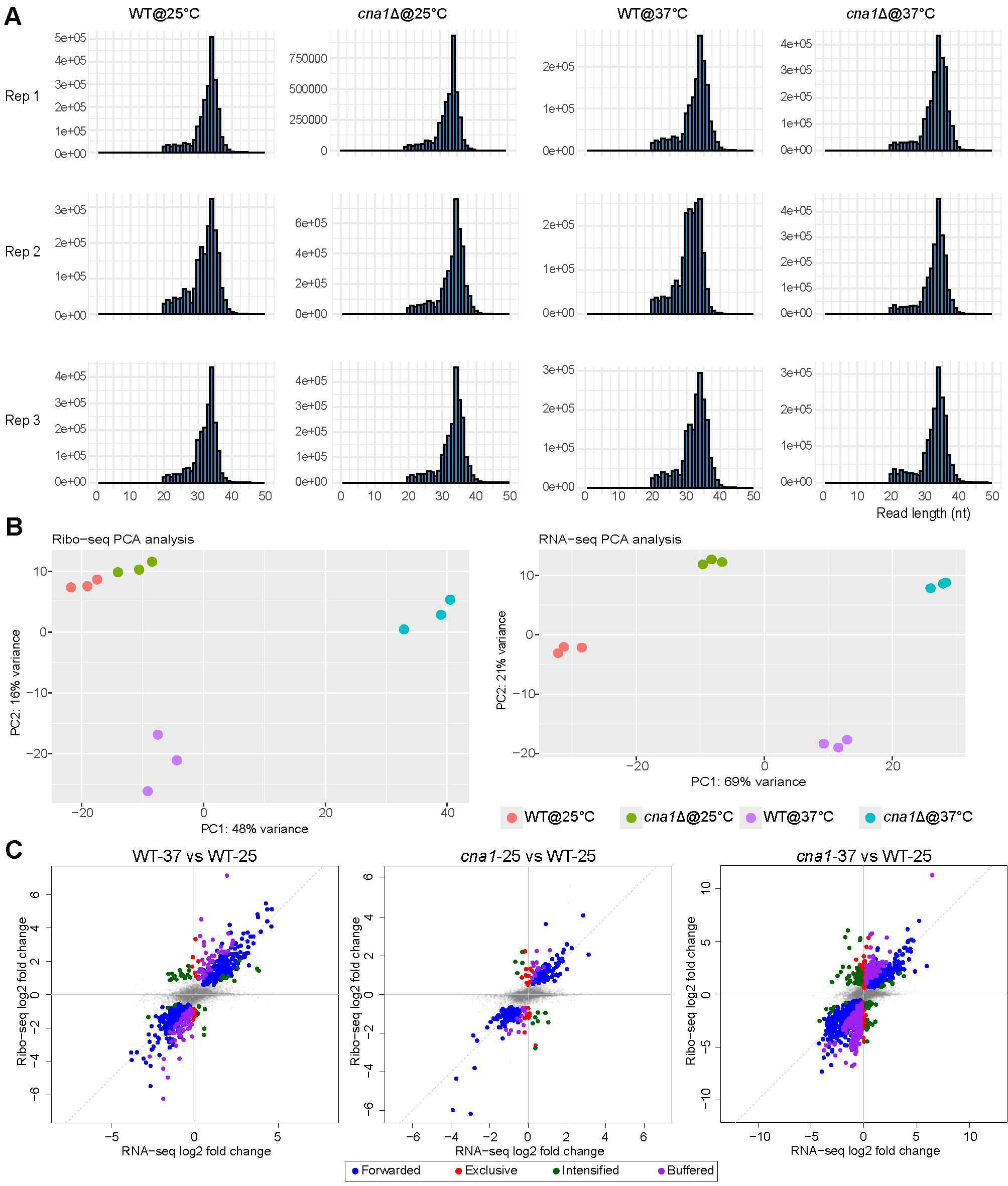

### Supplementary Figure S6

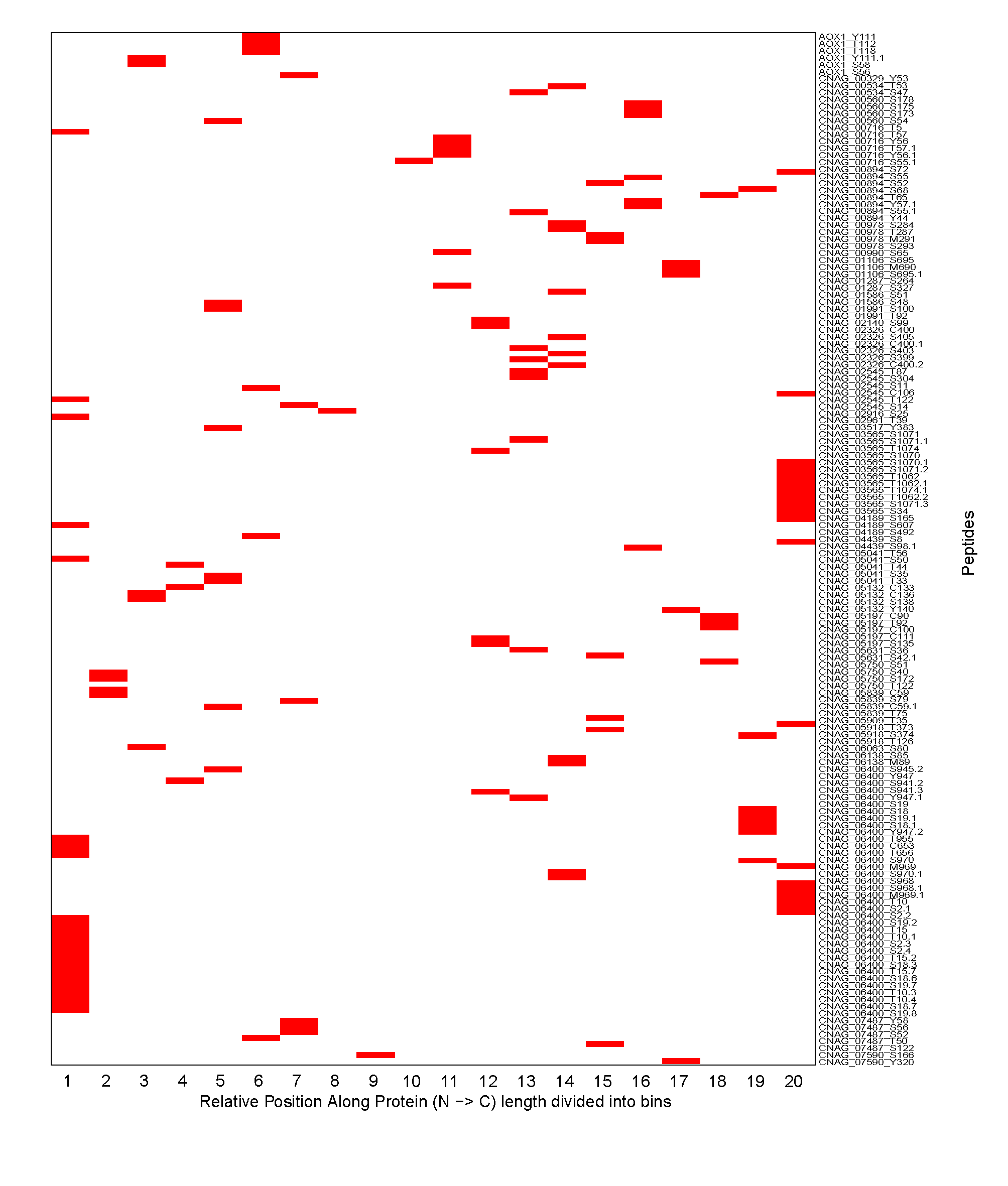

### Supplementary Figure S7

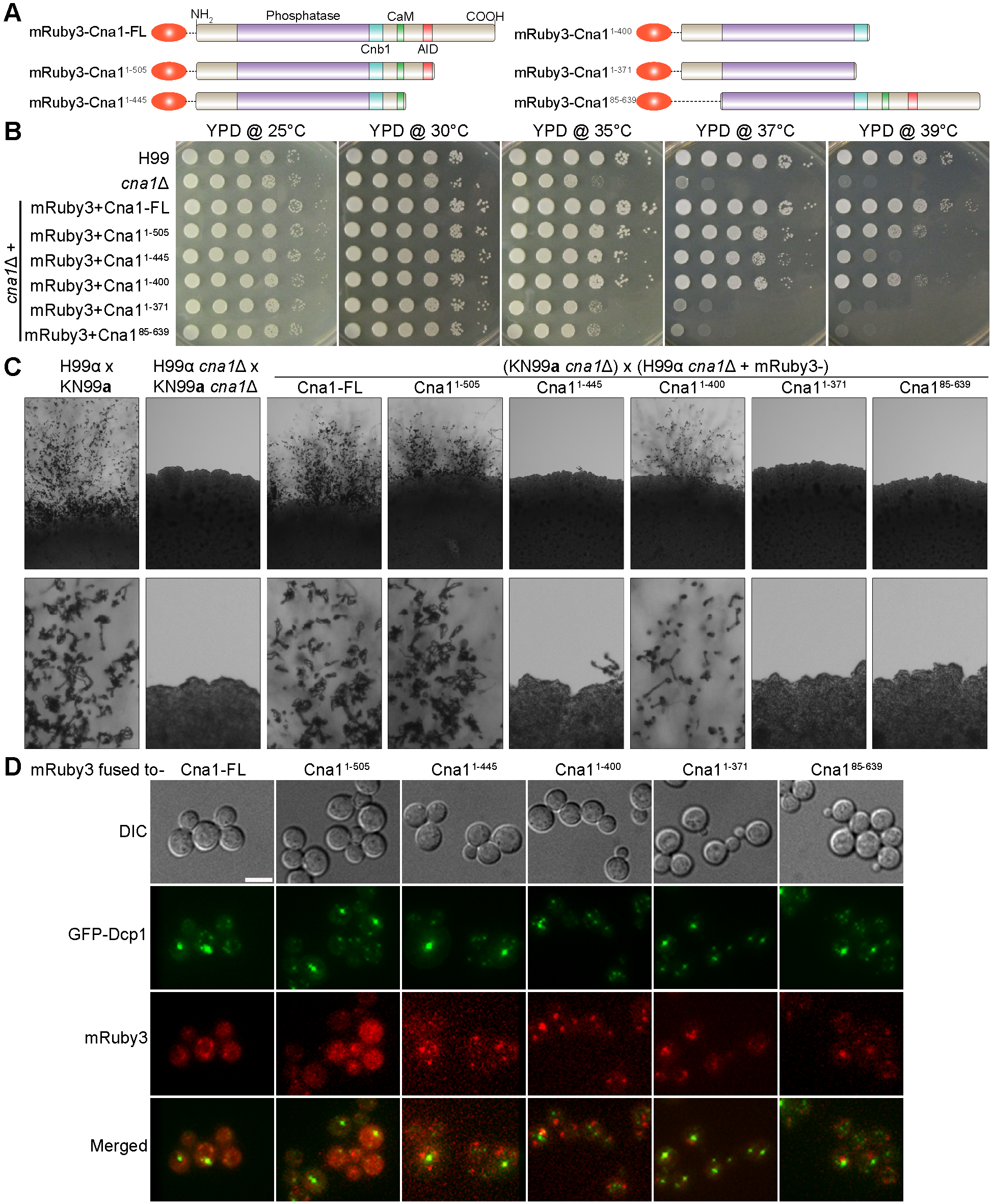

### Supplementary Figure S8

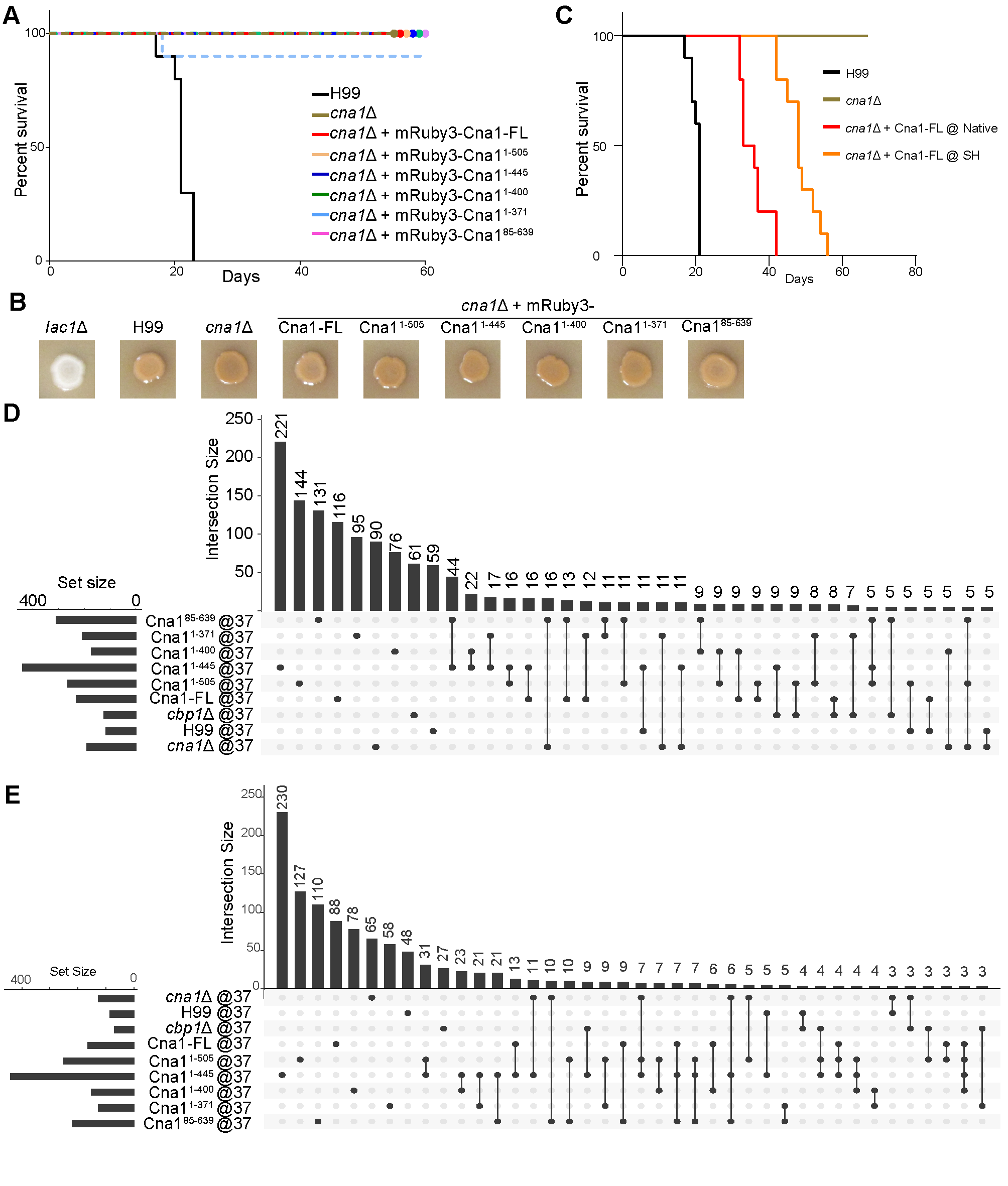
